# Maintenance of cell type-specific connectivity and circuit function requires Tao kinase

**DOI:** 10.1101/694265

**Authors:** Federico Marcello Tenedini, Maria Sáez González, Chun Hu, Lisa Hedegaard Pedersen, Mabel Matamala Petruzzi, Bettina Spitzweck, Denan Wang, Melanie Richter, Meike Petersen, Emanuela Szpotowicz, Michaela Schweizer, Stephan Sigrist, Froylan Calderon de Anda, Peter Soba

**Affiliations:** Neuronal Patterning and Connectivity laboratory, Center for Molecular Neurobiology (ZMNH), University Medical Center Hamburg-Eppendorf, Falkenried 94, 20251 Hamburg, Germany.; Neuronal Development laboratory, Center for Molecular Neurobiology (ZMNH), University Medical Center Hamburg-Eppendorf, Falkenried 94, 20251 Hamburg, Germany.; Electron microscopy unit, Center for Molecular Neurobiology (ZMNH), University Medical Center Hamburg-Eppendorf, Falkenried 94, 20251 Hamburg, Germany.; Institute of Biology, Free University Berlin, Takustr. 6, 14195 Berlin, Germany.

## Abstract

Sensory circuits are typically established during early development, yet how circuit specificity and function are maintained during organismal growth has not been elucidated. To gain insight we quantitatively investigated synaptic growth and connectivity in the *Drosophila* nociceptive network during larval development. We show that connectivity between primary nociceptors and their downstream neurons scales with animal size. We further identified the conserved Ste20-like kinase Tao as a negative regulator of synaptic growth required for maintenance of circuit specificity and connectivity. Loss of Tao kinase resulted in exuberant postsynaptic specializations and aberrant connectivity during larval growth. Using functional imaging and behavioral analysis we show that loss of Tao-induced ectopic synapses with inappropriate partner neurons are functional and alter behavioral responses in a connection-specific manner. Our data show that fine-tuning of synaptic growth by Tao kinase is required for maintaining specificity and behavioral output of the neuronal network during animal growth.

## Introduction

The function of a neuronal circuit is determined by synaptic strength and patterns of connectivity that allow information to flow in a specific manner to elicit behavior. Many circuits are formed during early development and undergo plastic changes including pruning and activity dependent refinement to establish and adjust functional connectivity ^1–5^. While the mechanisms of circuit establishment and refinement have been extensively studied in many systems, a less well understood process is how circuits remain functional while the animal and nervous system are still growing. This process requires scaling growth, adjustment of synaptic strength or both to maintain functional output despite changes in input resistance due to larger dendritic trees or muscles. In principal, circuit output in a growing animal could be maintained by homeostatic control of neurotransmitter release, postsynaptic receptor expression or by addition of synapses. While the former have been studied extensively by challenging synaptic function ^2^, the molecular mechanisms of how neuronal networks scale proportionally during animal growth and maintain their specificity and behavioral output are not well understood.

*Drosophila* larvae are an excellent system to study growth-related adjustments of circuit anatomy and function: the animals dramatically increase in size and enlarge their body surface 100-fold while maintaining structural and functional connectivity of their approximately 10,000 neurons ^6–8^. Both, the peripheral and central nervous system anatomically scale with animal growth: prominently, sensory dendrites of larval dendritic arborization (da) neurons cover the entire body wall and scale with the animal to maintain coverage ^9,10^. Similarly, synapse numbers and firing properties of motor neurons at the neuromuscular junction (NMJ) adjust during larval growth to maintain functional output ^11–14^. In the central nervous system (CNS), motor neuron dendrites proportionally increase their size during larval growth while maintaining the overall shape and receptive field domain ^8^.

Similar to the pioneering work on the *C. elegans* connectome, recent efforts to map Drosophila larval connectivity have now provided insight into circuit architecture and function of a more complex connectome ^15–19^. This includes the nociceptive class IV da (C4da) sensory neurons, which connect to an extensive downstream network and mediate responses to noxious mechanical and thermal stimulation, resulting in stereotyped rolling escape behavior ^20–22^. Recent electron microscopy (EM)-based reconstruction of the C4da neuron 2^nd^ order network revealed at least 13 subtypes consisting of 5 different local, 3 regional, 1 descending and 4 ascending classes of interneurons ^6^. In addition, this study has established that topography and sensory input are preserved in the early and late stage larval brain suggesting anatomical and functional scaling of the nociceptive network. Indeed, most larval behaviors including nociceptive responses are conserved throughout all stages suggesting that the majority of larval circuits maintain their function during animal growth ^23^. In recent years, a subset of C4da 2^nd^ order neurons has been studied in greater detail including A08n, DnB, Basin and mCSI neurons, which have been shown to be sufficient for nociceptive rolling behavior when activated by opto- or thermogenetic means ^18,24–28^. Functional network analyses by these and additional studies have revealed a hierarchical network organization, multisensory integration, and modality and position-specific network functions suggesting extensive processing and modulation of nociceptive inputs ^18,24,29^. This system thus offers a unique opportunity to probe how CNS circuit growth is regulated while preserving specific connectivity and functional output.

We and others have previously characterized A08n interneurons, which are major postsynaptic partners of C4da neurons required for nociceptive behavior ^24,27,28^. Here we characterize the developmental changes of the C4da-A08n circuit during larval growth at the synaptic level. We show that the number of pre- and postsynaptic sites as well as connectivity is proportionally increasing during larval development. We identified the conserved Ste20-like kinase Tao as a negative regulator of postsynaptic growth in A08n neurons. Loss of Tao function induces aberrant growth of dendrites and increased numbers of postsynaptic specializations. Strikingly, a subset of A08n postsynapses were no longer confined to the C4da presynaptic domain, but formed synapses with sensory neurons innervating adjacent regions of the neuropil. We show that these ectopic synapses are functional and result in altered network output and behavior. Our findings suggest that Tao kinase is required for maintenance of specific connectivity and function during animal growth by restricting postsynaptic growth in a circuit specific manner.

## Results

### Quantitative analysis of C4da and A08n neuron synapses

To evaluate the extent of synapses formed by neurons in the larval nociceptive circuit, we focused our efforts on establishing methods to visualize and quantify connections between C4da and A08n neurons, which display extensive synaptic contact along the entire ventral nerve cord (VNC) ^24^. To this end, we used three independent methods to assess synaptic connectivity by i) employing synapse-specific GFP reconstitution across synaptic partners (Syb-GRASP ^30^), ii) measuring the apposition of pre- and postsynaptic marker proteins ^31^ and iii) performing immuno-electron microscopy (EM) of synaptic markers labeling C4da-A08n neuron synapses ^24^. We first quantified the number of synaptic GRASP puncta from C4da-A08n neuron synapses in 3^rd^ instar larvae at 96 h after egg laying (AEL) using blind analysis of deconvolved 3D image stacks with automatic thresholding of synaptic puncta (details in methods). We consistently detected an average of 70-80 Syb-GRASP puncta per hemisegment (Fig. 1A-C, F).

**Figure 1:**
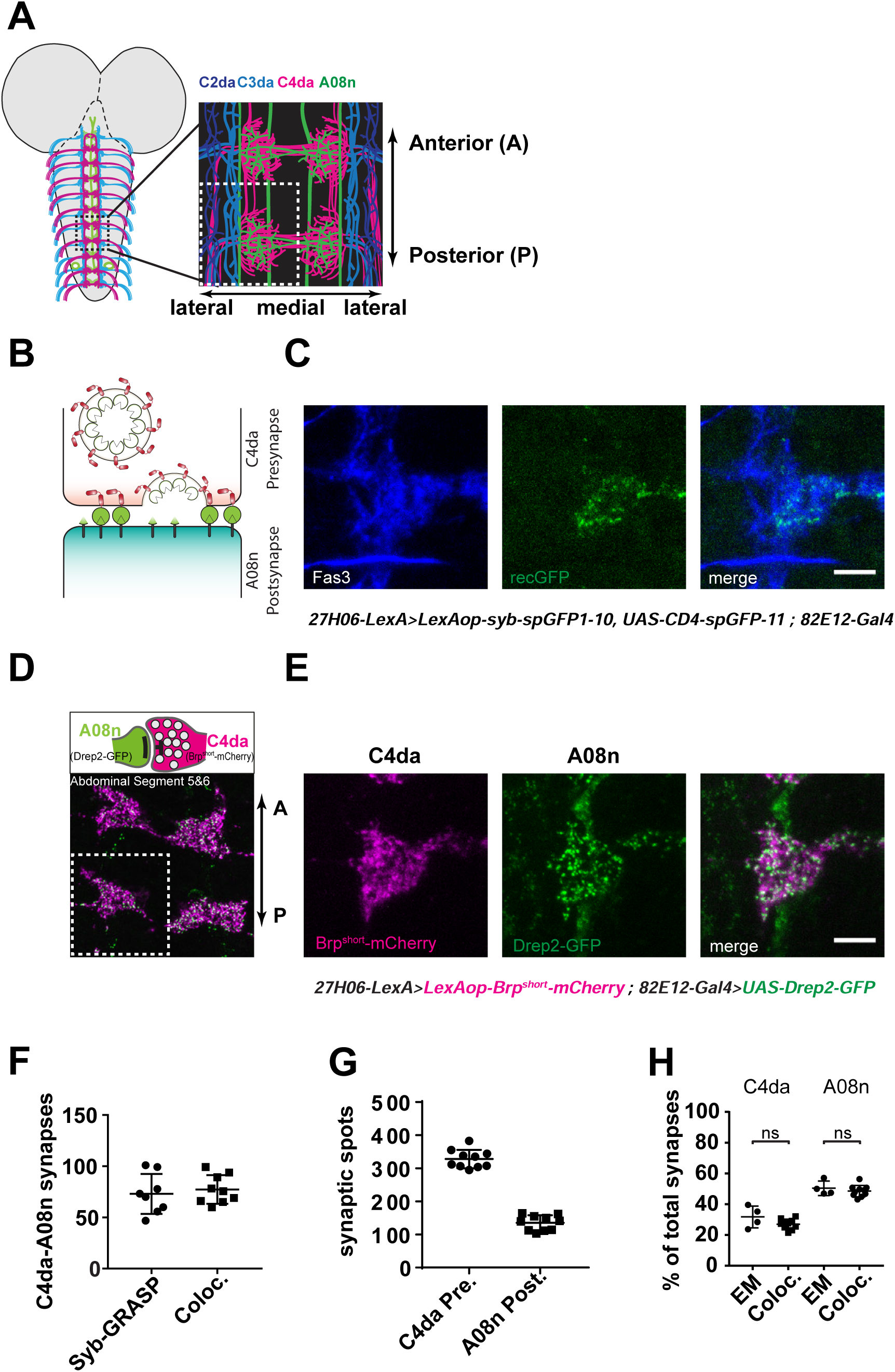
Quantitative analysis of C4da-A08n neuron connectivity. (A) Schematic model of a larval brain with A08n neurons (green), C4da (magenta) and C2da/C3da axon projections (blue/cyan) into the VNC and schematic expansion of abdominal segments 5 and 6. (B) Schematic model of Syb-GRASP mediated GFP reconstitution. (C) C4da-A08n Syb-GRASP signal in a single hemisegment showing anti-Fas3 staining (blue), reconstituted GFP (green) and merge. Scale bar = 5 µm. (D) C4da pre- (Brp^short^-mCherry) and A08n postsynaptic (Drep2-GFP) marker expression and example confocal image of abdominal segments 5 and 6 showing expressed markers in C4da (magenta) and A08n (green). (E) Hemisegment showing Brp^short^-mCherry in C4da (magenta), Drep2-GFP in A08n (green) and merge. Scale bar = 5 µm. (F) Quantification of Syb-GRASP puncta compared to synaptic marker colocalization. Syb-GRASP n = 8, colocalization n = 9. ±SD (G) Quantification of C4da pre- and A08n postsynaptic marker puncta per hemisegment. n = 10. ±SD. (H) Comparison of EM and light microscopic marker colocalization analyses. Percentages of counted C4da presynapses connected to A08n or A08n postsynapses connected to C4da showed no significant differences between both methods. EM: C4da n=4, A08n n=4, confocal: C4da n=10, A08n n=10. C4da: *P*= 0.06, A08n: *P*=0.43. ±SD. Unpaired two-tailed *t*-test.

To facilitate comparison of GRASP synapse numbers with C4da and A08n neuron synaptic sites, we used the active zone marker Brp^short^-mCherry ^32^ to label C4da neuron-specific presynapses. In order to label A08n postsynaptic densities, we used Drep2-GFP previously shown to discretely label postsynaptic densities when expressed in mushroom body Kenyon cells ^33^ (Fig. 1D,E). We detected close apposition of Brp^short^-mCherry and Drep2-GFP at discrete foci in areas of C4da-A08n contact, and analyzed the number of co-localized C4da-A08n neuron synaptic puncta using automatic thresholding of apposed Brp/Drep2 puncta together with a distance threshold similar to previous work ^31,34^ (Fig. 1F, Supplementary Figure 1A-C, see methods for details). Synapse numbers determined using this approach were comparable to numbers from our Syb-GRASP analysis, suggesting that both methods allowed us to estimate C4da-A08n neuron connectivity. We further analyzed the number of C4da presynaptic and A08n postsynaptic puncta in different abdominal segments: overall numbers were similar from segment to segment, with C4da neurons displaying about 2-3-fold higher presynaptic counts compared to A08 postsynapses (Supplementary Figure 1A-C). Moreover, C4da-A08n neuron synapse counts correlated more with the number of A08n post-than C4da presynaptic sites (Supplementary Figure 1D,E).

Lastly, we performed immuno-EM labeling of C4da-A08n connectivity in larvae expressing Brp^short^-mCherry (C4da) and Drep2-GFP (A08n). We first counted the total number of synapses on C4da and A08n neurons and then calculated the percentage of synapses that contained A08n postsynapses and C4da presynapses, respectively (Fig. 1H, Supplementary Figure 1F). We found that ∼30% of C4da neuron active zones formed synapses with A08n neurons and that ∼50% of Drep2-GFP labeled A08n postsynaptic sites contacted C4da presynapses. Strikingly, the relative C4da-A08n synapse numbers we observed using EM and light microscopy were indistinguishable. Taken together, our analysis shows that light microscopic pre- and postsynaptic marker apposition and Syb-GRASP analysis provide valid representations of C4da-A08n neuron connectivity.

### C4da-A08n neuron connectivity scales with larval growth

We next wanted to assess C4da and A08n neuron connectivity across larval development. *Drosophila* larvae grow extensively after hatching and dendrite length and synaptic numbers at the NMJ and in the CNS have been shown to increase for the subsets of neurons investigated so far ^6,8,10,34,35^. We analyzed C4da-A08n neuron synapse numbers from 48 h to 120 h AEL using Syb-GRASP and synaptic marker apposition (Fig. 2A,B). Both methods showed a comparable linear increase of synaptic numbers from 48 h to 96 h AEL, with synapse numbers close to doubling every 24 h. We observed a decline of C4da-A08n synapses at 120 h AEL using Syb-GRASP but not colocalization analysis, hinting at potential changes in their connectivity in wandering stage larvae (Fig. 2A). C4da neuron presynaptic puncta kept increasing until 120 h, while A08n postsynaptic counts plateaued at 96 h (Fig. 2C,D). We then calculated the ratio of C4da-A08n neuron connections across development and found that the relative C4da presynaptic output to A08n neurons displayed mild alterations between the analyzed developmental timepoints, but remained within a range between 20-30% (Fig. 2E). In contrast, we observed a significant increase in synapse/postsynapse ratios for A08n neurons from 48 h to 72 h AEL indicating a developmental increase in their relative connectivity to C4da neurons during the transition from 2^nd^ to 3^rd^ instar stages (Fig. 2E). Taken together, these data show that C4da-A08n neuron synaptic numbers scale with larval growth and undergo stage-specific adjustments in connectivity.

**Figure 2:**
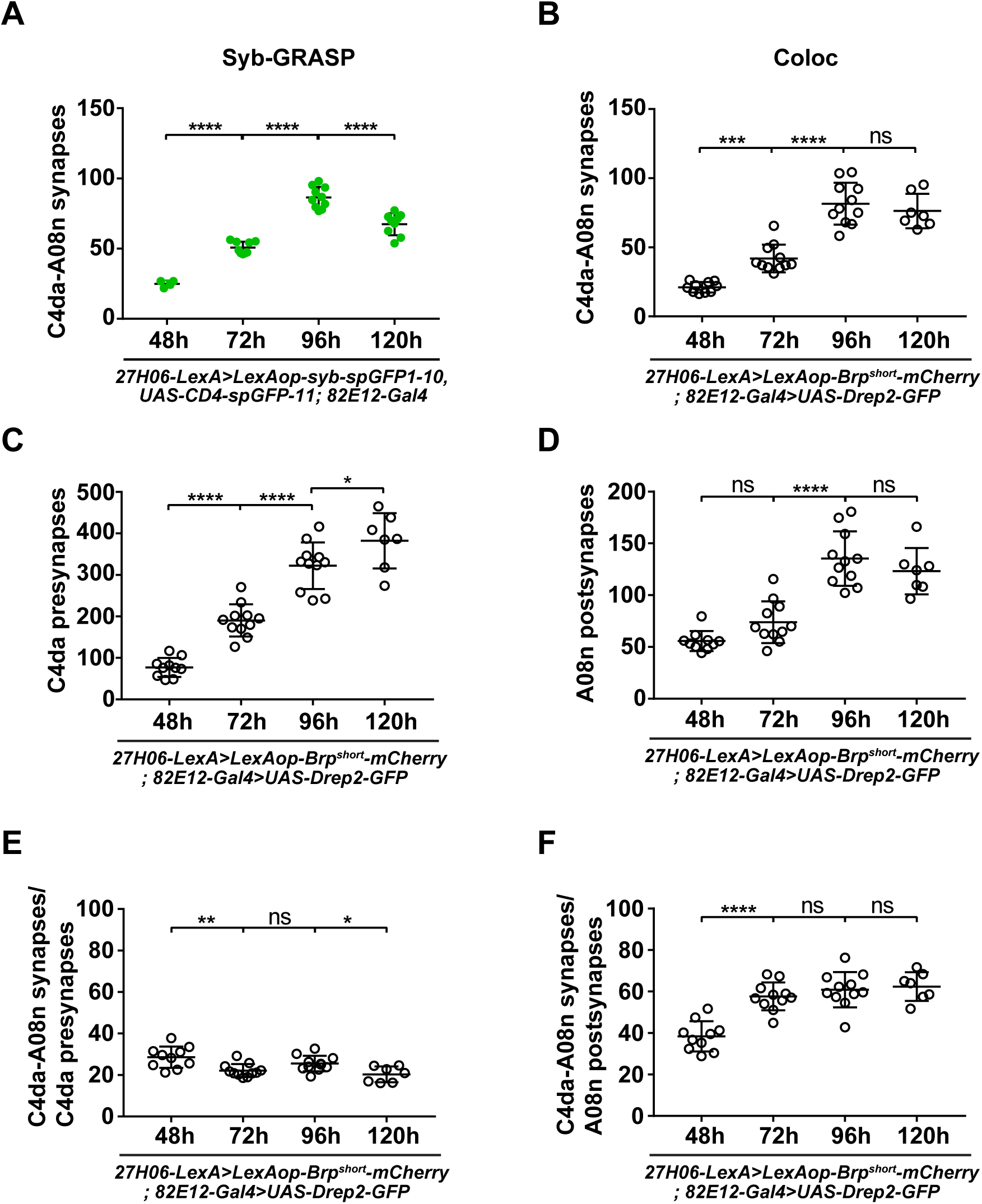
Scalar increase of C4da-A08n connectivity during larval growth. (A) Quantification of C4da-A08n synapses during development from 48h AEL until 120h AEL with Syb-GRASP. 48h: n=4, 72h: n=10, 96h: n=10, 120h: n=9. **** *P* < 0.0001. ANOVA with multiple comparisons and Sidak’s *post-hoc* test (for exact *P* values and statistics see Supplementary Table 1). (B-D) Quantification of C4da-A08n synaptic connectivity during development from 48h AEL until 120h AEL using colocalization of synaptic markers. Quantification of (B) C4da-A08n synapses, *** *P* < 0.001, **** *P* < 0.0001. (C) C4da presynapses and (D) A08n postsynapses. (E) C4da-A08 synapse to C4da presynapse ratios (in percent) during development from 48 AEL until 120h AEL. (F) C4da-A08 synapse to A08n postsynapse ratios (in percent) during development from 48 AEL until 120h AEL. 48h: n=10, 72h: n=11, 96h: n=11, 120h: n=7, * *P* < 0.05, ** *P* < 0.01, *** *P* < 0.001, **** *P* < 0.0001. ±SD, ANOVA with multiple comparisons and Sidak’s *post-hoc* test (for exact *P* values and statistics see Supplementary Table 1).

### Tao kinase function restricts postsynaptic growth of A08n neurons

We next focused on how A08n postsynaptic growth might control synaptogenesis with C4da neurons. In a candidate RNAi screen for growth-related genes we identified Tao kinase as a regulator of synaptic growth in A08n neurons. We perturbed Tao function in A08n or C4da neurons using RNAi-mediated knockdown (*Tao^RNAi^*) or by overexpression of a hyperactive form of Tao (Tao^CA^) ^36^ and analyzed synapse numbers using our newly established methods. A08n-specific knockdown of Tao resulted in a significant increase of A08n postsynaptic puncta at 96 h AEL (Fig. 3A,B’). In contrast, Tao hyperactivation caused a reduction of Drep2-GFP puncta. A08n neuron expression of *Tao^RNAi^* did not significantly affect C4da presynaptic or C4da-A08n synaptic numbers, while Tao^CA^ overexpression strongly reduced both, suggesting that hyperactivation of Tao function negatively regulates C4da-A08n neuron synaptic connectivity (Fig. 3A, B-B’’). We sought to validate these results using Syb-GRASP and found that while *Tao^RNAi^* in A08n neurons did not affect C4da-A08n synapse numbers, Tao^CA^ expression reduced GRASP puncta to a comparable extent as observed by our co-localization analysis (Fig. 3C,D). We also tested if Tao kinase was involved in presynaptic control of C4da-A08n neuron connectivity. Interestingly, C4da neuron-specific *Tao^RNAi^* expression did not affect synaptic marker numbers at 96 h AEL, while Tao^CA^ overexpression strongly reduced C4da pre-, A08n post- and C4da-A08n synaptic numbers (Supplementary Figure 2A,B-B’’). These data suggest that presynaptic Tao kinase hyperactivation has a trans-synaptic effect, while postsynaptic reduction of Tao levels affects A08n postsynaptic growth independent of C4da neurons.

**Figure 3:**
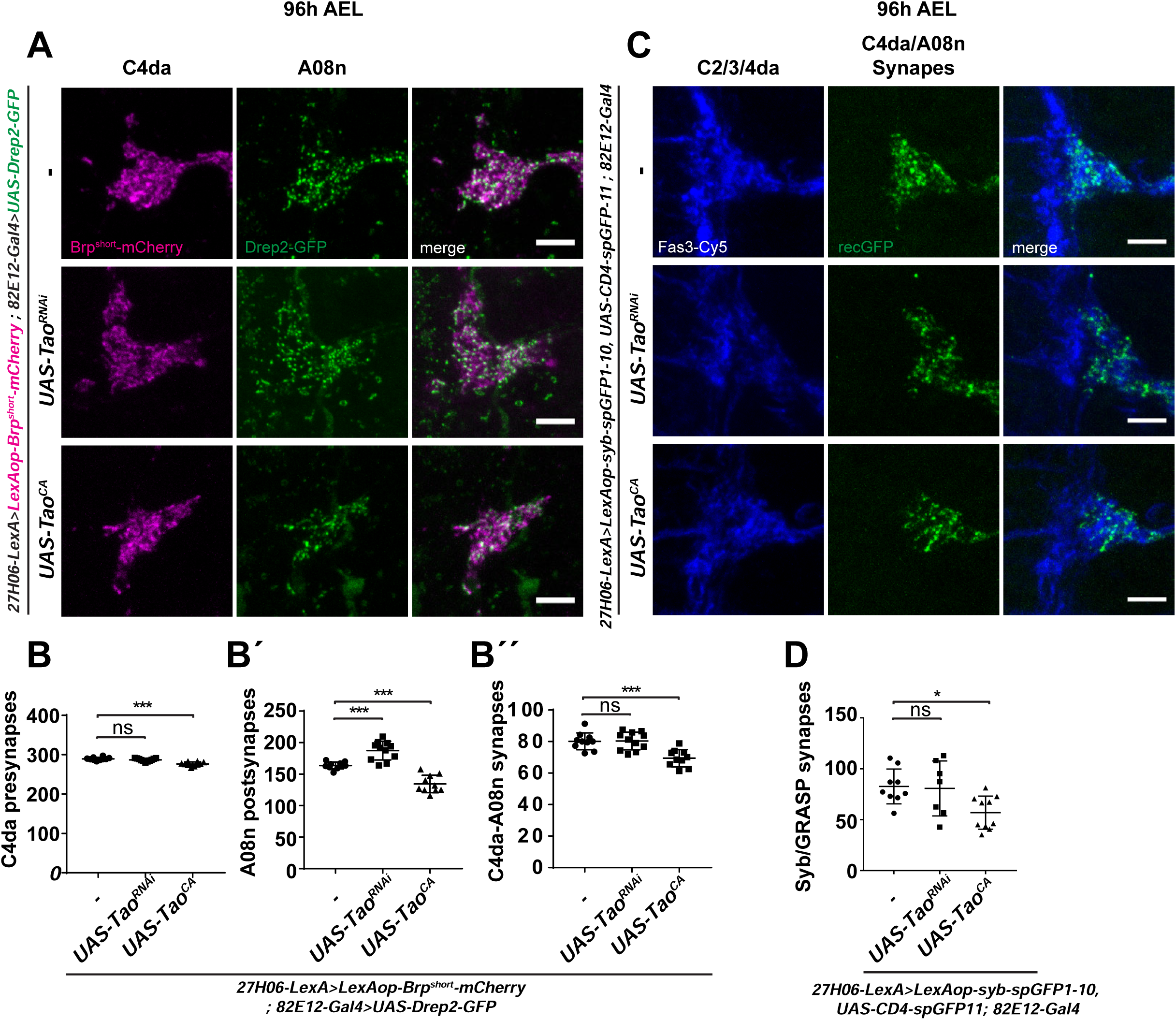
Tao kinase regulates postsynaptic growth of A08n neurons. (A) Confocal images of hemisegments in control or with *Tao^RNAi^* and Tao^CA^ expression in A08n neurons using synaptic markers labeling of C4da presynapses (magenta) and A08n postsynapses (green). Scale bar = 5 µm. (B) Quantification of C4da pre-, (B’) A08n post- and (B’’) colocalized C4da-A08n synaptic markers in control or with *Tao^RNAi^* and Tao^CA^ expression in A08n neurons. \*\*\**P<*0.001, **** *P* < 0.0001. ±SD, ANOVA with multiple comparisons and Dunnett’s *post-hoc* test (for exact *P* values and statistics see Supplementary Table 1). *Control* n = 10, *UAS-Tao^RNAi^* n = 11, *UAS-Tao^CA^* n = 10. (C) Confocal images of Syb-GRASP-labeled C4da-A08n synapses. Hemisegments of control animals or with Tao^RNAi^ and Tao^CA^ expression in A08n neurons together with anti-Fas3 staining are shown. Fas3 labels C2da, C3da and C4da sensory axons (blue) overlapping with reconstituted GFP signal within the C4da neuron domain (green). Scale bar = 5 µm. (D) Quantification of C4da-A08n neuron synapses using Syb-GRASP under control conditions or with *Tao^RNAi^* and Tao^CA^ expression in A08n neurons. *Control* n = 9, *UAS-Tao^RNAi^* n = 7, *UAS-Tao^CA^* n = 10. \**P<*0.05. ±SD, ANOVA with multiple comparisons and Dunnett’s *post-hoc* test (for exact *P* values and statistics see Supplementary Table 1).

As *Tao^RNAi^* in A08n neurons resulted in an increase of postsynaptic Drep2-GFP puncta, we further analyzed the localization of the presumptive additional postsynaptic compartments. We expressed Drep2-GFP together with a morphological marker (CD4-tdTomato) in A08n neurons while perturbing Tao function (Supplementary Figure 2C,C’,D-D’’). Reduction of Tao activity using Tao^RNAi^ resulted in striking dendritic overgrowth and concomitant increase in postsynaptic puncta of A08n neurons. Immunostaining with an anti-Fas3 antibody, which specifically labels C2da, C3da and C4da sensory axons, revealed that A08n dendrites and postsynapses extended into the adjacent domains of C2da and C3da neurons, which align laterally to the medial triangular-shaped C4da axon projections. Conversely, hyperactivation of Tao kinase in A08n neurons resulted in a reduced dendritic field and fewer postsynapses. Neither perturbation affected the number of A08n postsynapses per dendritic volume suggesting that Tao activity co-regulates dendritic and synaptic growth (Supplementary Figure 2D-D’’).

We compared loss of Tao-induced synaptic and dendritic growth changes in A08n neurons with overexpression of constitutively active Ras (*UAS-Ras85D^V12^*) or Rac1 (*UAS-Rac1^V12^*), which were previously shown to promote synaptic growth at the fly NMJ ^37,38^. Strikingly, Ras^V12^ but not Rac1^V12^ overexpression phenocopied the loss of Tao (Supplementary Figure 3A, B-B’’) indicating that Tao acts in a Ras-like manner to coordinate dendritic and synaptic growth. In both cases, A08n neurons displayed a comparable increase of postsynapses and dendritic volume with unchanged density (Supplementary Figure 3B’’). In contrast, expression of constitutive active Rac1 led to a strongly altered dendritic field with loss of volume and postsynapses, additionally resulting in lowered postsynaptic site densities. Collectively, these data show that Tao kinase function in A08n neurons negatively co-regulates dendritic growth and postsynaptic numbers, thus limiting synaptic input to the C4da neuron presynaptic domain.

### Loss of Tao kinase function promotes ectopic growth throughout development

We then analyzed the impact of loss of Tao kinase function on C4da-A08n neuron synaptic markers during larval development. *Tao^RNAi^* in A08n neurons did not strongly affect C4da presynapse numbers compared to controls except at 72h AEL (Fig. 4A, Supplementary Figure 4A-D). In contrast, A08n postsynaptic numbers remained continuously elevated after loss of Tao and, remarkably, kept increasing at 120 h AEL (Fig.4B). Consistently, C4da-A08n neuron synapse numbers were significantly elevated at 48 h and 72 h, and particularly at 120 h AEL (Fig. 4C). These experiments suggest that Tao function is required throughout development to restrict A08n postsynaptic numbers and in part also C4da-A08n neuron synapses. Loss of Tao function increased the synapse/presynapse ratio in C4da neurons at most time points suggesting an overall shift in C4da neuron connectivity towards A08n neurons (Fig. 4D). In contrast, synapse/postsynapse ratios in A08n were decreased at 72 h and 96h AEL indicating a relative increase in alternative presynaptic inputs of A08n neurons (Fig. 4E). These results are consistent with the observed dendritic overgrowth phenotype with A08n dendrites invading adjacent neuropil domains upon loss of Tao (see Supplementary Figure 2C, D).

**Figure 4:**
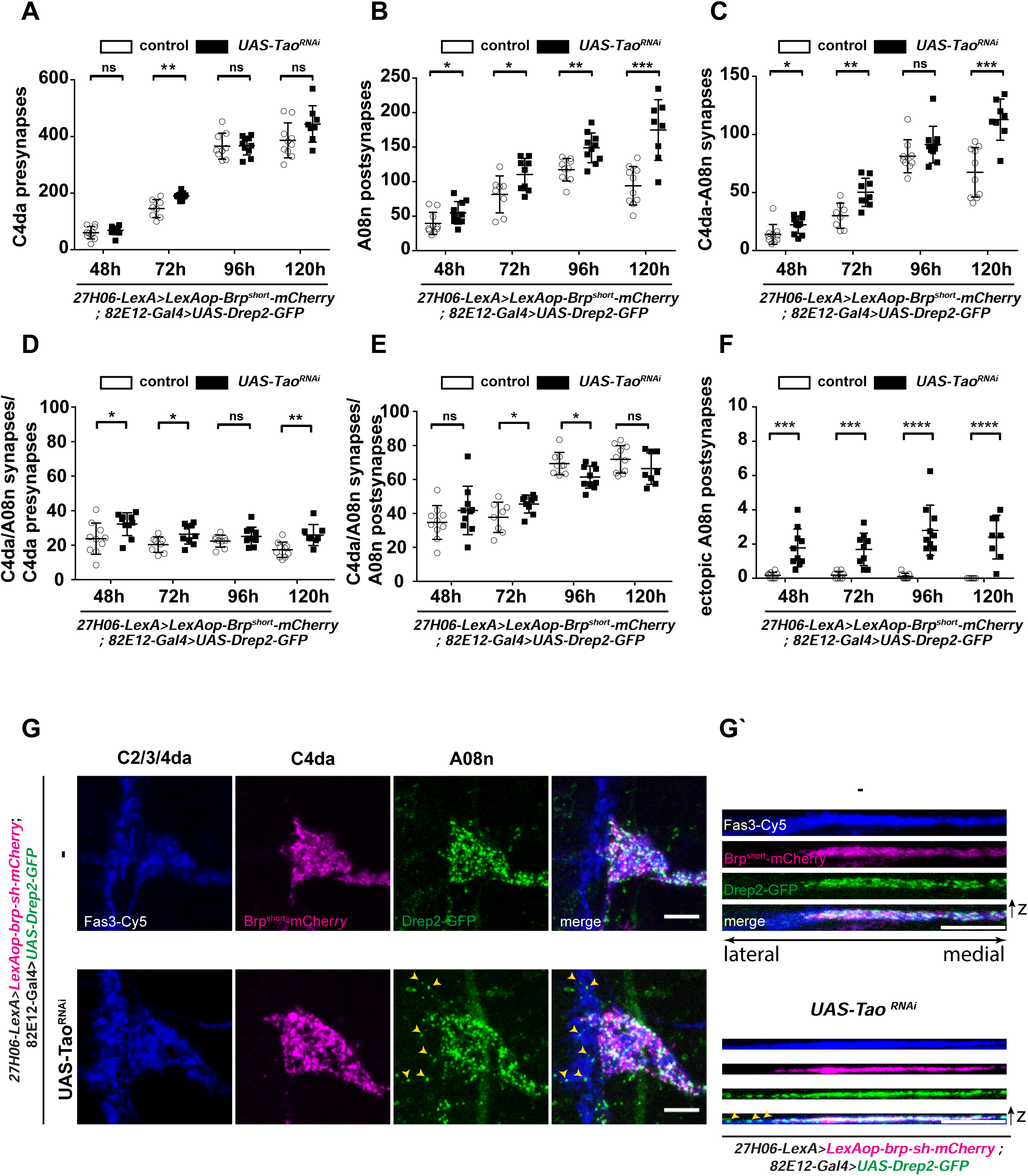
Tao kinase restricts postsynaptic growth and connectivity of A08n neurons during larval development. (A-C) Quantitative analysis of C4da-A08 neurons synaptic profiles from 48h AEL to 120h AEL in control or with *Tao^RNAi^* expression in A08n neurons. (A) C4da neuron presynapse (B) A08n postsynapses and (C) C4da-A08n synapse numbers. \**P<*0.05, \*\**P<*0.005, \*\*\**P<*0.001. ±SD, unpaired two-tailed *t*-test (for exact *P* values and statistics see Supplementary Table 1). *Control* 48h n = 10, 72h n = 8, 96h n = 9, 120h n = 10. *UAS-Tao^RNAi^* 48h n = 10, 72h n = 9, 96h n = 10, 120h n = 8. (D) C4da-A08 synapse to C4da presynapse ratios (in percent) during development from 48 AEL to120h AEL in control or with *Tao^RNAi^* expression in A08n neurons. \**P<*0.05, \*\**P<*0.005. ±SD, unpaired two-tailed *t*-test (for exact *P* values and statistics see Supplementary Table 1). *Control* 48h n = 10, 72h n = 8, 96h n = 9, 120h n = 10. *UAS-Tao^RNAi^* 48h n = 10, 72h n = 9, 96h n = 10, 120h n = 8. (E) C4da-A08n synapse to A08n postsynapse ratios (in percent) during development from 48 AEL to 120h AEL in *control* or with *Tao^RNAi^* expression in A08n neurons. \**P<*0.05, **** *P* < 0.0001. ±SD, unpaired two-tailed *t*-test (for exact *P* values and statistics see Supplementary Table 1). Control 48h n = 10, 72h n = 8, 96h n = 9, 120h n = 10. UAS-Tao^RNAi^ 48h n = 10, 72h n = 9, 96h n = 10, 120h n = 8. (F) Quantification of ectopic A08n postsynapses in C2da/C3da domain during development from 48h AEL until 120h AEL. \*\*\**P<*0.001, **** *P* < 0.0001. ±SD, unpaired two-tailed *t*-test (for exact *P* values and statistics see Supplementary Table 1). Control 48h n = 10, 72h n = 8, 96h n = 9, 120h n = 10. *UAS-Tao^RNAi^* 48h n = 10, 72h n = 9, 96h n = 10, 120h n = 8. (G) Confocal images of hemisegments in *control* or with *Tao^RNAi^* expression in A08n neurons. Synaptic markers labeling C4da presynapses (magenta), A08n postsynapses (green) and anti-Fas3 immunolabeling C2da, C3da and C4da sensory axons (blue) at 96h AEL. Yellow arrowheads show ectopic A08n postsynapses within the C2da/C3da domain under *Tao^RNAi^* conditions. (G’) XZ projections of each channel in (G) are shown. Scale bar = 5 µm.

We next examined the developmental profile of ectopic postsynaptic puncta of A08n neurons, which were not localized within the C4da neuron presynaptic domain upon loss of *Tao* function. We therefore analyzed the number of postsynaptic Drep2-GFP puncta that overlapped with the C2da/C3da presynaptic domain labeled by anti-Fas3 immunostaining (Fig. 4F,G,G’). In controls, we rarely observed A08n postsynapses localized outside of the C4da neuron presynaptic domain. In contrast, *Tao^RNAi^* in A08n neurons led to ectopic A08n postsynapses that were displaced laterally within the adjacent domain of C2da/C3da sensory neuron projections. Ectopic A08n postsynapses were already present at 48 h AEL and persisted to a similar degree throughout development (Fig. 4F). This suggests that Tao kinase function is required to prevent ectopic postsynaptic sites by restricting the A08n postsynaptic domain.

### Conserved Tao kinase activity regulates postsynaptic growth

Overexpression of hyperactive Tao kinase resulted in a strong decrease of A08n Drep2-GFP puncta (see Fig. 3), which might indicate kinase activity-dependent regulation of postsynaptic growth in A08n neurons. To test this hypothesis further and to probe potentially conserved Tao activity, we asked if the closest human orthologue, Tao kinase 2 (hTaoK2), was capable of compensating for the loss of *Drosophila* Tao. TaoK2 has recently been shown to affect dendritic and synaptic development in mammals and has been linked to Autism spectrum disorders (ASDs) based on patient mutations that alter its kinase activity ^39–41^. We compared the ability of hTaoK2 or a kinase activity-impaired ASD-linked variant (hTaoK2^A135P^) to rescue loss of Tao in A08n neurons in respect to dendritic morphogenesis and synaptic overgrowth (Fig 5A, Supplementary Figure 5). Quantitative analysis of A08n dendrites revealed that loss of Tao in A08n neurons resulted in an increase in the number and length of dendrite branches invading the lateral C2/3da domain of the neuropil. hTaoK2 but not hTaoK2^A135P^ restored A08n dendritic branching to control levels and was able to fully suppress *Tao^RNAi^*–induced lateral branches (Supplementary Figure 5A-D). Similarly, we found that hTaok2 overexpression was able to fully rescue *Tao^RNAi^*-induced A08n postsynaptic overgrowth and prevent formation of lateral ectopic postsynapses (Fig. 5B-E). In contrast, kinase-impaired hTaok2^A135P^ displayed attenuated rescue activity: although it could partially normalize A08n postsynaptic and C4da-A08n synapse numbers, ectopic Drep2-GFP puncta and dendrites were still present. These results show that Tao and hTaok2 are functionally conserved and that its kinase activity is important to restrict dendritic and ectopic postsynaptic growth in A08n neurons.

**Figure 5:**
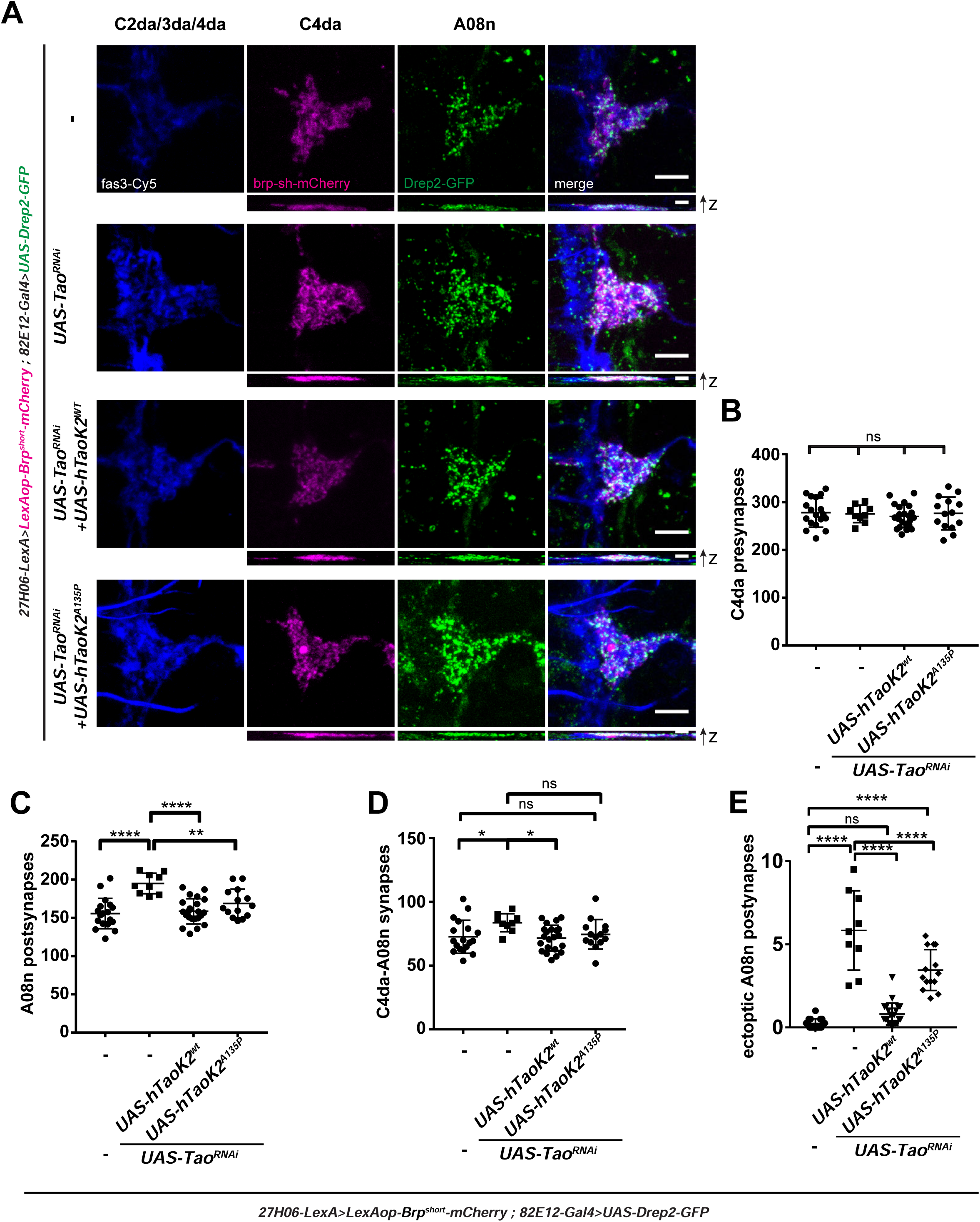
Conserved Tao kinase activity regulates postsynaptic growth and connectivity of A08n neurons. (A) Confocal Images of larval VNC hemisegments (96h AEL) in control or with *Tao^RNAi^* expression in A08n neurons, without or with co-expression of hTaoK2^wt^ or hTaoK2^A135P^. Images show anti-Fas3 immunostaining labeling C2da, C3da and C4da sensory axons (blue), with synaptic marker expression labeling C4da presynapses (magenta) and A08n postsynapses (green). Scale bar = 5 µm. XZ projections of each channel are shown below. Scale bar = 2 µm. (B-E) Quantitative analysis of synaptic profiles in control or with Tao^RNAi^ expression in A08n neurons, without or with co-expression of hTaoK2^wt^ or hTaoK2^A135P^ for (B) C4da neuron presynapses, (C) A08n postsynapses, (D) C4da-A08n synapses and (E) ectopic A08n postsynapses within the C2da/C3da domain. * *P* < 0.05, ** *P* < 0.01, **** *P* < 0.0001. ±SD, ANOVA with multiple comparisons and Dunnett’s *post-hoc* test (for exact *P* values and statistics see Supplementary Table 1). *Control*: n = 18, *UAS-Tao^RNAi^*: n = 9, *UAS-Tao^RNAi^ + hTaoK2^wt^*: n = 22, *UAS-Tao^RNAi^ + hTaoK2^A135P^*: n = 14.

### Tao loss of function in A08n neurons generates aberrant functional connectivity

We next addressed if loss of Tao-induced ectopic A08n postsynaptic structures were indeed forming functional synapses. Axons of C2da, C3da and C4da somatosensory neurons form laminated non-overlapping structures in the VNC, with C4da neurons displaying the most medial projections followed by C3da and C2da neurons ^42^. Based on the lateral displacement of A08n neuron postsynaptic sites after Tao loss of function, we hypothesized that C3da neurons might be a major subset of ectopic presynaptic partners. To assess if C3da and A08n neurons indeed form synaptic connections, we performed Syb-GRASP experiments across larval development with and without perturbation of *Tao* function in A08n neurons. Using a line expressing Syb-spGFP1-10 in C3da and chordotonal (cho) neurons (*nompC-LexA*), few GRASP puncta were detected between C3da and A08n neurons in controls from 24 to 120h AEL, consistent with the observed confinement of A08n dendrites to the medial C4da axonal domain (Fig. 6A,B, Supplementary Figure 6A). In contrast, *Tao^RNAi^* in A08n neurons resulted in a strong increase in C3da-A08n synaptic GRASP puncta, first detectable at 48 h AEL. No Syb-GRASP puncta were detected between A08n and the more dorso-laterally positioned cho neuron presynapses suggesting that Tao function is required during larval development to restrict C3da-A08n synapse formation.

**Figure 6:**
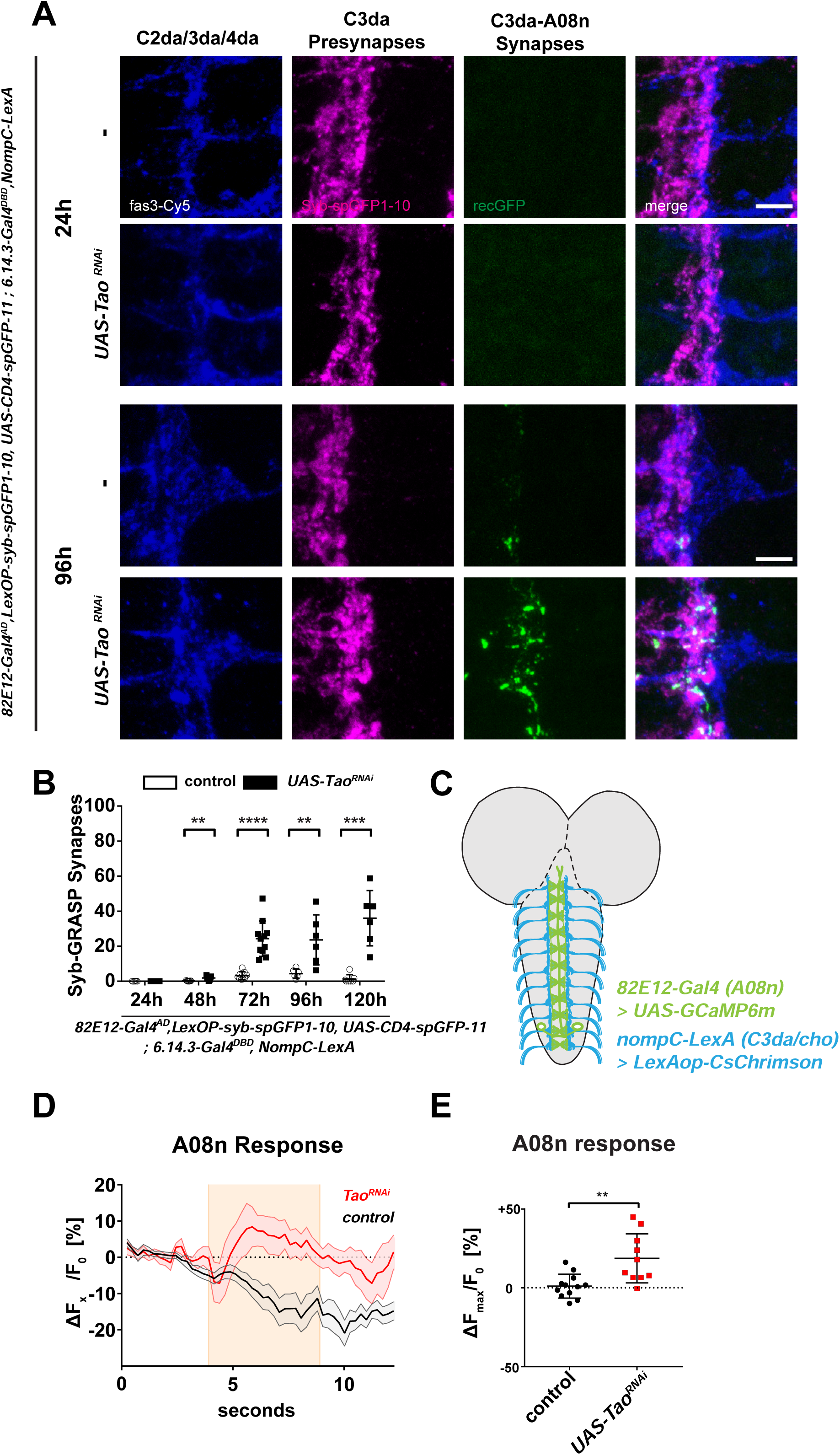
A08n neurons form functional ectopic synapses with C3da neurons after loss of Tao. (A) Confocal images of Syb-GRASP-labeled C3da-A08n synapses (24 and 96h AEL). Representative images of larval VNC hemisegments in control or with Tao^RNAi^ expression in A08n neurons showing anti-Fas3 labeling of C2da, C3da and C4da sensory axons (blue), presynaptic spGFP1-10 expressed in C3da (magenta) and reconstituted GFP signal marking C3da-A08n Synapses (green). Scale bar = 5 µm. (B) Quantification of C3da-A08n Syb-GRASP synapses in control or with *Tao^RNAi^* expression in A08n neurons. ** *P* < 0.01, *** *P* < 0.001, **** *P* < 0.0001, 24h *P=* ns, 48h *P=* 0.0017, 72h *P= <*0.0001, 96h *P=* 0.0294, 120h *P=* 0.0007. ±SD, unpaired two-tailed *t*-test. 24h control n = 5, *UAS-Tao^RNAi^* n = 6, 48h control n=7, *UAS-Tao^RNAi^* 1.893 n=7, 72h control: n=9, *UAS-Tao^RNAi^*: n=11, 96h control n=6, *UAS-Tao^RNAi^* n=6, 120h control n=7, *UAS-Tao^RNAi^* n=6. (C) Schematic larval brain showing A08n neurons (green) and C3da sensory dendrite VNC projections (blue) and indicating expression of *UAS-GCaMP6m* in A08n and *LexAop-CsChrimson* in C3da/cho. (D) Calcium responses of GcaMP6m-expressing A08n neurons after optogenetic activation of C3da/cho neurons using CsChrimson (5s, 630 nm, indicated by shaded area), with or without *Tao^RNAi^* expression in A08n neurons. Data show mean change in percent [(ΔF/F0)-1, (±SEM indicated by shaded regions]. *Control* n = 12, *UAS-Tao^RNAi^* n = 10. (E) Quantification of maximum A08n responses to C3da activation in percent [(ΔFMax/F0)-1)] comparing control and *Tao^RNAi^* expression in A08n neurons. \*\**P* < 0.005, *P=* 0.0024. ±SD, unpaired two-tailed *t*-test. *Control* n=12, *UAS-Tao^RNAi^* n=10.

To further test if C3da-A08n synapses were functional under these conditions, we performed *in vivo* calcium imaging experiments. We activated C3da/cho neurons (*nompC-LexA*) using the red-shifted optogenetic actuator CsChrimson ^43^ and monitored calcium signals in A08n neurons with or without Tao perturbation using the calcium-responsive fluorescent protein GCaMP6m (Fig. 6C-E). Under control conditions, C3da/cho neuron activation did not drive calcium responses in A08n (Fig. 6D,E). In contrast, activation of C3da/cho neurons in larvae expressing *Tao^RNAi^* in A08n neurons reproducibly resulted in A08n calcium responses, demonstrating that ectopic C3da and A08n synapses are functional.

We also tested if loss of Tao affects functional connectivity of C4da and A08n neurons. Using optogenetic activation of C4da neurons, we detected a significant decrease in A08n neuron responses after loss of Tao compared to controls (Supplementary Figure 6B-D). These data show that loss of Tao in A08n neurons gives rise to functional ectopic connectivity with C3da sensory neurons while partially impairing C4da-A08n neuron physiological output.

### Ectopic Tao-dependent connectivity alters somatosensory network function and behavioral action selection

To dissect the impact of Tao-dependent connectivity changes, we analyzed behavioral consequences of Tao loss of function in this system. We focused on C3da, cho and C4da-dependent behaviors based on their converging circuits and functional role in noxious responses. C3da neurons are primarily involved in innocuous touch and noxious cold responses, which result in stop and turn behavior or full body contraction, respectively ^44–46^. Similarly, cho neurons respond to noxious cold and high-frequency vibration giving rise to very similar behaviors including contraction/hunching ^47,48^. Moreover, C3da and cho neurons contribute to nociceptive rolling behavior in response to noxious mechanical stimulation or vibration-induced co-activation, respectively ^18,24^. We first tested if *Tao^RNAi^* in A08n neurons caused mechanonociception defects and if Tao kinase activity was required (Fig. 7A, Supplementary Figure 7A). Expression of *Tao^RNAi^* using an A08n-specific split-Gal4 line resulted in reduced mechanonociceptive responses, which could be fully rescued by overexpression of hTaok2 but not its kinase-impaired hTaok2^A135P^ variant. Comparable results were obtained using optogenetic activation of C4da neurons (Supplementary Figure 7B). However, synaptic output of A08n neurons was not severely affected, as CsChrimson-mediated activation of A08n neurons with or without *Tao^RNAi^* resulted in comparable nociceptive rolling responses (Supplementary Figure 7C). These results suggest that C4da-A08n synaptic transmission is partially impaired due to Tao manipulation, consistent with reduced A08n responses after optogenetic C4da neuron activation (see Supplementary Figure 6B-D).

**Figure 7:**
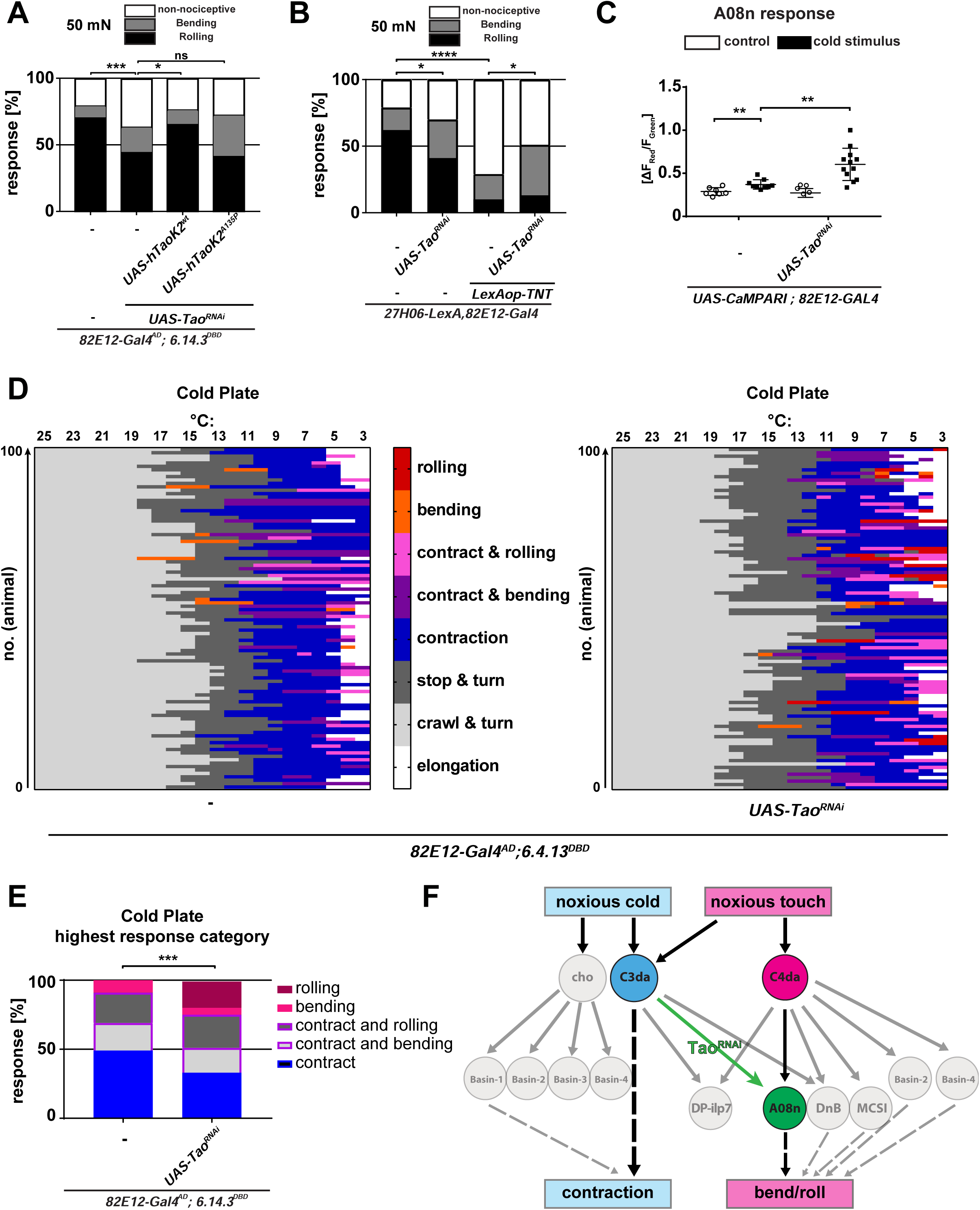
Loss of Tao-induced A08n connectivity changes alter behavioral action selection. (A) Mechanonociceptive behavioral response of third instar larvae (96h AEL) in *control* or with *Tao^RNAi^* without or with co-expression of hTaoK2^wt^ or hTaoK2^A135P^ in A08n neurons. Responses to the second mechanical stimulation with a 50 mN *von Frey* filament are shown (Nociceptive rolling and bending or non-nociceptive responses). \**P* < 0.05, \*\*\**P* < 0.001. *Control*: n = 98, *UAS-Tao^RNAi^* n = 99, *UAS-Tao^RNAi^ + hTaoK2^wt^*: n = 99, *UAS-Tao^RNAi^ + hTaoK2^A135P^*: n = 98. *Control* vs *UAS-Tao^RNAi^*: *P =* 0.0006, *UAS-Tao^RNAi^* vs *UAS-Tao^RNAi^ +hTaok2^wt^*: *P =* 0.0109, *UAS-Tao^RNAi^* vs *UAS-Tao^RNAi^ +hTaok2^A135P^*: *P =* 0.1006, χ^2^ test. (B) Mechanonociceptive behavioral response of third instar larvae (96h AEL) in control or *Tao^RNAi^*, Tetanus toxin light chain (TNT), or *Tao^RNAi^* with TNTe expression in A08n neurons. Responses to the second mechanical stimulation with a 50 mN *von Frey* filament are shown. \**P* < 0.05, \*\*\*\**P* < 0.00005. *Control* n = 100, *UAS-Tao^RNAi^*: n = 100, *LexAop-TNTe*:n = 100, *UAS-Tao^RNAi^ + LexAop-TNTe*: n = 100. *Control* vs *UAS-Tao^RNAi^*: *P =* 0.0249, *Control* vs *LexAop-TNT*: *P =* >0.0001, *LexAop-TNT* vs *UAS-Tao^RNAi^ + LexAop-TNT*: *P =* 0.0201, χ^2^ test. (C) A08n calcium responses to a 4°C cold stimulus based on CaMPARI photoconversion without or with Tao^RNAi^ expression in A08n neurons. \*\**P*<0.005, control vs control cold *P=* 0.0066, control cold vs UAS-Tao^RNAi^ cold *P=* 0.002. Unpaired two-tailed *t*-test. *Control* n=7, *Control* cold: n=9, *UAS-Tao^RNAi^* control n = 9, *UAS-Tao^RNAi^* cold: n=12. (D) Cold plate assay ethogram showing individual larval behaviors during continuous temperature reduction (25-3°C) without or with *Tao^RNAi^* in A08n neurons (stop & turn, contraction, contraction & bending, contraction & rolling, bending and rolling). *Control* n = 100. *UAS-Tao^RNAi^* n = 100. (E) Highest response categories of larvae shown in (D). \*\*\**P =* 0.0001, χ^2^ test. (F) Model of C4da and C3da/cho circuits and Tao-dependent network changes. Ectopic connectivity between C3da and A08n neurons after loss of Tao results in a shift in noxious cold behavior towards C4da/A08n-dependent rolling.

To address if Tao-dependent ectopic C3da-A08n neuron connectivity contributed to mechanonociceptive behavior, we expressed Tetanus toxin light chain (TNT) in C4da neurons while reducing Tao function in A08n neurons. Interestingly, nociceptive behavior was partially restored compared to TNT expression alone, possibly due to functional activation of A08n neurons by C3da neurons (Fig. 7B). In contrast, C4da neuron specific optogenetic activation revealed that A08n neuron output is required for Tao-dependent behavioral changes, as simultaneous silencing blocked any differences (Supplementary Figure 7D). These findings suggest a direct contribution of loss of Tao-induced C3da-A08n neuron connections to mechanonociceptive behavior.

Lastly, we examined whether ectopic connectivity between C3da and A08n neurons could affect noxious cold responses, which are mediated by C3da/cho neurons ^45,47^. We first assayed if A08n neurons could respond to a cold stimulus after loss of Tao. To this end, we analyzed calcium-dependent photoconversion of the calcium integrator CaMPARI ^49^ in A08n neurons in response to noxious cold. We observed a significant increase in CaMPARI photoconversion in Tao^RNAi^-expressing A08n neurons following a noxious cold stimulus, while controls displayed only low level responses (Fig. 7C). Next we addressed if this altered activation of A08n neurons could affect noxious cold behavior using a cold plate assay. Freely crawling larvae (96h AEL) were cooled from 25 °C to 3 °C within 50 seconds, which resulted in sequential stop & turn behavior (mean onset at 14,79 °C) followed by contraction (mean onset at 9,8 °C), bending or mixed behavior (contract & bend, contract & roll) in control animals (Fig. 7D,E). However, loss of Tao in A08n neurons resulted in a significant number of animals displaying pure rolling behavior thus altering action selection at noxious cold temperatures (<10°C). Loss of Tao in A08n neurons also shifted the onset of stop & turn behavior and bending to higher or lower temperatures, respectively (Supplementary Figure 7E). Taken together, these data show that loss of Tao-induced connectivity changes between C3da and A08n neurons functionally alter noxious C3da/cho-dependent cold and C4da-dependent mechanonociceptive responses thus perturbing modality-specific behavioral action selection.

## Discussion

Evolution shaped precise neuronal networks that elicit appropriate sensory modality-dependent actions. To select for accurate behavioral responses or sequences, combinations of feedback or feedforward inhibition, disinhibition and co-activation are thought to result in command-like decisions selecting a certain behavior or induce probabilistic behavioral sequences ^18,48,50–52^. Yet organismal size increase during juvenile development also requires proportional growth of its nervous system to maintain connectivity and thus the appropriate behavioral output. For example, zebrafish larvae already display prey catch behavior 5 days post fertilization, which is maintained and refined until adulthood despite undergoing 5-fold growth in body length and further nervous system development ^53^. Similarly, *Drosophila* larvae display preservation of most behaviors including nociceptive and mechanosensory responses from early to late stages ^23^ suggesting functional conservation of most circuits throughout larval growth. To investigate and understand the underlying principles of circuit maintenance that ensure consistent patterns of behavior during animal growth we combined quantitative synapse analysis, behavioral and physiological assays within the larval nociceptive network. Our analysis of C4da sensory and A08n neuron connectivity across larval development showed that synaptic contacts are linearly increasing up to 4-fold from 48-96 h. Interestingly, C4da neurons maintain a connectivity ratio of around 25-30% with A08n throughout larval life, suggestive of tight control of synapse addition to maintain not only partner specificity, but likely also balanced innervation and output to all postsynaptic targets. Our findings are consistent with EM and light microscopic data of C4da neuron connectivity and axonal structure at early and late larval stages^6,54^. Topography and connectivity of C4da neurons and the nociceptive network were shown to be preserved, with dendritic cable length and synaptic connections increasing 4-5 fold during larval growth. Similarly, larval visual circuits and NMJs show growth-specific adjustments in synaptic numbers as well ^7,11,14,34,55^. Notably, C4da neurons form polyadic synapses clustering 3-4 postsynaptic sites of different partners at most active zones, and these structures remain stable over development ^6^. While our light microscopic methods cannot resolve polyadic synapses, our analyses nonetheless faithfully depict the relative increase in C4da-A08n neuron connectivity, in strong agreement with the EM analyses. Moreover, our data show that the A08n postsynaptic connectivity ratio with C4da neurons increases from 48h to 72h suggesting a developmental adjustment in the A08n circuit. Scalar growth of neurons has also been described for subsets of retinal cells in other systems including fish and mammals. Goldfish type 1 and 2, mouse alpha- and cat alpha-/beta-type ganglion cells establish their dendritic coverage early in development and expand their territory several-fold during postnatal development in sync with the increasing size of the retina^56–59^. This suggests that similar and possibly conserved mechanisms govern proportional growth of networks to maintain function during organismal size increase.

The observed proportional growth of receptive fields is required to maintain spacing and connectivity, yet the underlying molecular mechanisms have not been fully elucidated. Both, activity-dependent and -independent mechanisms, are known to contribute to circuit refinement and stability ^5^. How either of these is required for network scaling and specificity during organismal growth remains to be elucidated in detail. Our results point to growth-related factors as major players in this process: Tao function restricts dendritic and synaptic growth during *Drosophila* larval development, which is critical for maintaining circuit specificity. Tao prevents dendritic overgrowth in a cell-autonomous fashion suggesting its activity is tightly controlled, possibly by extrinsic signals required to coordinate synaptic growth of partner neurons. This suggests a prime role for Tao kinase in maintaining the growth-related balance of dendrites, synapses and network function. Previous studies have implicated that dendrite placement and growth of dendritic arbors can alter or increase synaptic input, respectively ^8,60,61^. Similarly, synaptic domains are frequently defined by exclusion mechanisms suggesting that overriding these restrictive cues can result in aberrant connectivity ^62^. While connection specificity seems to be in part encoded by cell type-specific expression of synaptic adhesion molecules, domain-specific axon targeting and dendrite placement are likely critical to prevent innervation of alternative neuronal targets. Consistently, driving neuronal growth by constitutive-active Ras overexpression resulted in Tao-like dendritic and postsynaptic overgrowth. In contrast, Rac1 overactivation had detrimental effects on A08n dendritic field organization and significantly reduced A08n postsynapse density. While Ras and Rac-driven signals have been implicated in dendritic and synaptic growth, they differentially regulate dendritic field organization and synaptogenesis ^63,64^. Thus the regulation of growth-promoting and -limiting factors like Tao kinase might play a pivotal role not only in scaling growth, but also in restricting synaptic connectivity to the appropriate domain.

Many other sensory networks including tectal neurons in tadpole and the larval visual circuit show extensive sensory input and activity-dependent control of dendritic arborization and synaptic growth during development ^34,55,65^. Similarly, steroid hormone signaling by ecdysone is required to scale motor neuron dendrites and activity during transition from the 2^nd^ to the 3^rd^ instar stage ^8^. While it remains to be shown if Tao kinase is involved in activity-dependent adjustments of dendritic and synaptic growth, sensory input and neuronal activity have been shown to adjust nociceptive output and fine-scale topography of C4da neuron axon projections, respectively ^27,54^. This suggests that the nociceptive circuit is subject to structural and functional plasticity responding to both growth and activity-related cues. As Tao kinase signaling has been linked to dendrite growth and cytoskeletal regulation ^36,40,66,67^, it is likely that it controls the intracellular machinery required to coordinate actin and microtubule dynamics in response to extrinsic signals, thus regulating scaled dendritic and synaptic growth.

Our data show that Tao’s role in proportional growth is likely conserved as its closest human orthologue Taok2 was able to substitute for *Drosophila* Tao function. Interestingly, an ASD-linked kinase-impaired Taok2 variant ^39^ did not recover function suggesting that Tao kinase activity is essential for its role in growth regulation. Moreover, it suggests that mutations altering Taok2 activity regulate connectivity during development, thus contributing to network changes linked to ASDs. Loss of Taok2 in mice results in altered dendritic morphology and spine stability ^40,41,67^, and Taok2 knockout mice display impaired social behavior and learning flexibility ^39^. Similarly, other ASD mouse models show altered mechanosensory circuit connectivity and behavior ^68^, which likely contribute to ASD-linked sensory alterations. Similarly to our Tao phenotype, loss of the growth pathway inhibitor PTEN in mice results in dendritic overgrowth and increased spine density, giving rise to ASD-like physiological and behavioral defects ^69^. Thus growth-related alterations in neuronal morphology can substantially alter connectivity and contribute to pathological behavioral changes.

Recent studies have described a hierarchical circuit architecture and multisensory integration in the larval somatosensory network, which integrates the activity of distinct sensory subsets and determines specific behaviors by inhibition or facilitation ^18,24,25,27,48^. While individual larval sensory neuron subsets have been shown to respond to specific modalities like vibration (cho)^70^, noxious cold (cho, C3da)^45,47^, gentle (C2da, C3da)^21,46^ and harsh touch (C4da)^21,71^, overlap in 2^nd^ order neuron innervation also results in extensive sensory integration (Fig. 7F). Vibration-induced cho neuron activation has been shown to boost C4da neuron-dependent rolling responses by integration of both inputs via basin 2^nd^ order neurons and increased output to command-like neurons (goro)^18^. Similarly, mechanosensitive C2da, C3da and C4da neurons are cooperatively required for mechanonociception by facilitating C4da and A08n neuron responses via integrating neuromodulatory feedback neurons (DP-ilp7)^24^. Thus tight regulation of these circuits ensures sensory modality-specific responses even during larval growth, which is key to appropriate actions important for survival. Here we demonstrate that under circumstances of aberrant growth of A08n neurons, connectivity can be altered and have a direct impact on behavior. In agreement with Tao being required for sensory circuit integrity, our analyses showed that loss of Tao in A08n neurons induced functional ectopic synapses with C3da neurons specifically during larval growth. C3da and cho neurons were shown to mediate noxious cold responses resulting in contraction behavior, which is predominantly selected over other behaviors including C4da and A08n neuron-dependent nociceptive rolling and bending ^21,24,45,47^. While C3da-A08n neuron connectivity is minimal in wild-type animals, loss of Tao-induced A08n neuron overgrowth induced exuberant C3da-A08 synapses and enhanced A08n neuron activation by noxious cold. As a result, cold-induced behavior was partially shifted towards A08n-mediated rolling suggesting a direct contribution of these ectopic synapses. Although the C3da neuron downstream network has not been fully elucidated, they do share network partners with C4da and cho neurons, which among others likely include Basin neurons ^18,25,48^. Similarly to C3da, activation of cho or basin-1 neurons can induce contraction behavior, while basin-2 activation results primarily in head bending, which suppresses contraction/hunching by feedforward inhibition ^48^. A similar inhibitory mechanisms might normally suppress rolling during C3da/cho neuron activation, thus selecting for contraction behavior. Tao-induced ectopic C3da-A08n connectivity might bypass this mechanism by direct activation of the rolling response (Fig. 7F). While exuberant C3da-A08n neuron connectivity likely does not explain all observed behavioral changes, it hints at how an imbalance in interconnected networks can alter behavioral action selection. Thus the growth-restricting activity of Tao prevents ectopic connections with alternative partners including C3da neurons. This ectopic connectivity with other sensory neurons, sensing different modalities, results in altered behavioral patterns and cross-modality responses. Based on the broad expression of Tao kinase family members in the nervous system of invertebrates and vertebrates, we anticipate Tao having a general function regulating proportional growth of circuits to maintain network specificity and behavior.

## Methods

### Drosophila melanogaster stocks

All stocks were maintained at 25 °C and 70% rel. humidity with a 12 h light/dark cycle on standard fly food unless noted otherwise. All transgenic lines were maintained in a white mutant (w^−^) or yellow white mutant (y^−^,w^−^) background. Stocks were obtained from the Bloomington (BDSC) Drosophila stock centers unless otherwise indicated. We used the following lines:

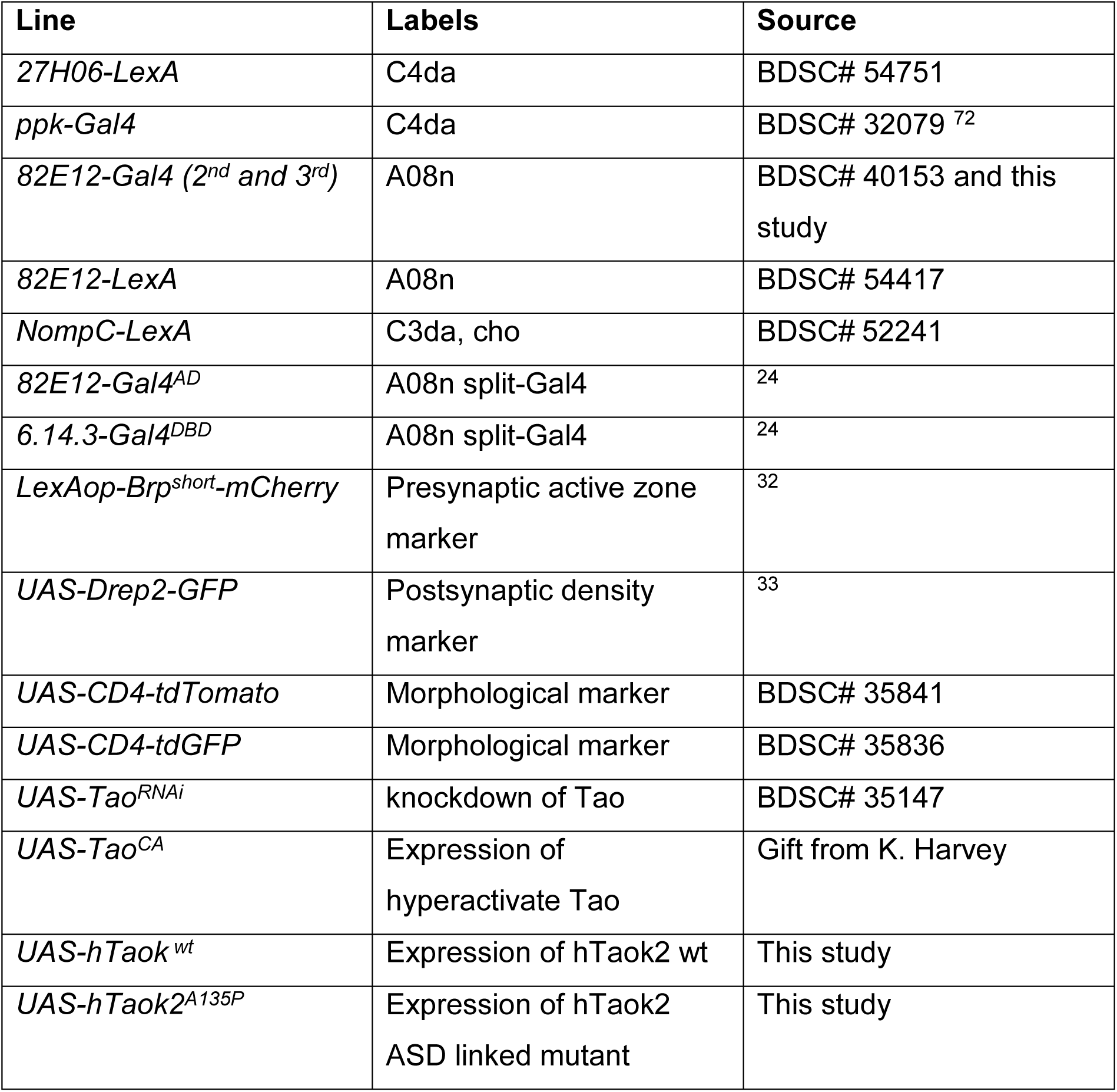
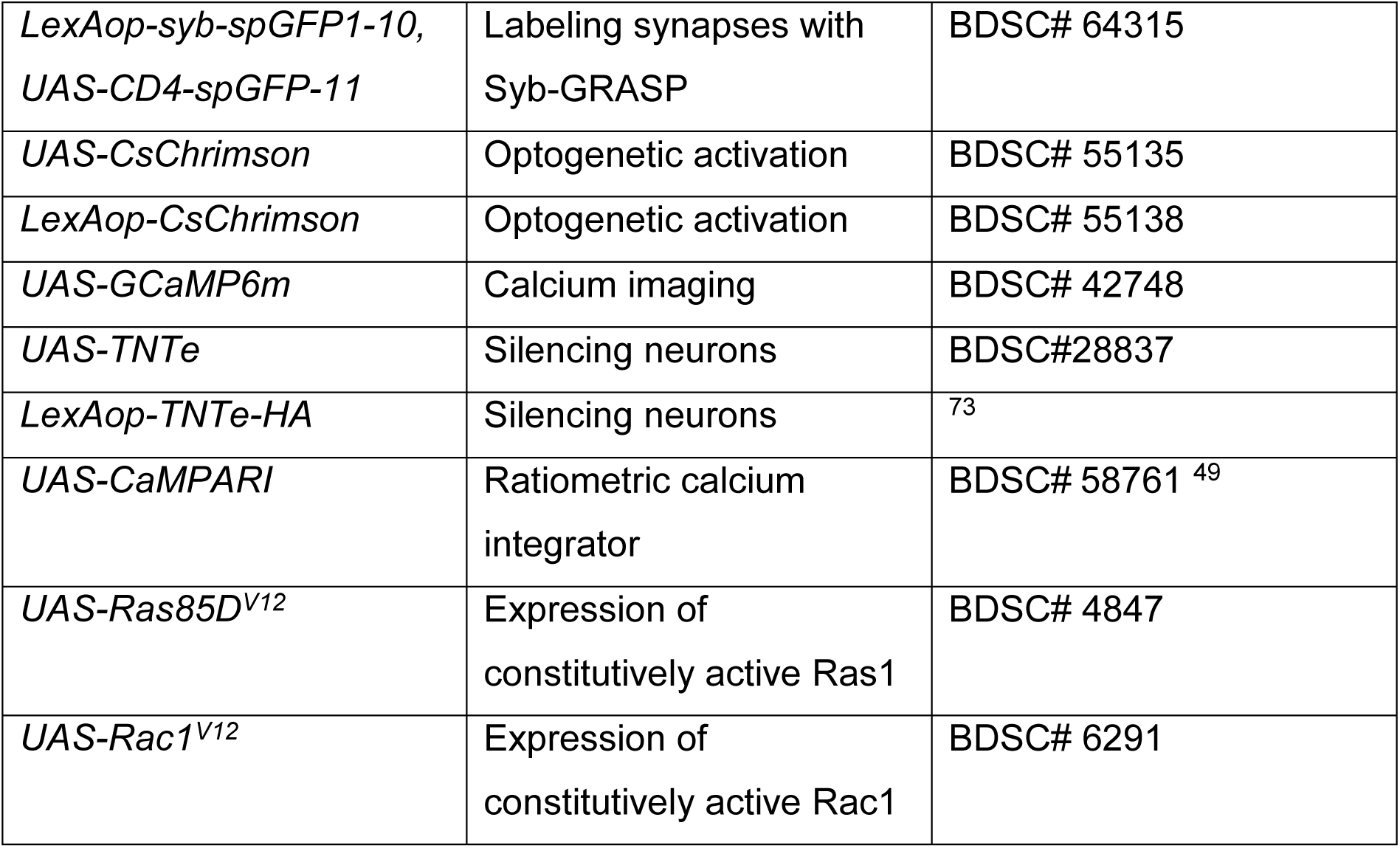

| Line | Labels | Source |
| --- | --- | --- |
| *27H06-LexA* | C4da | BDSC# 54751 |
| *ppk-Gal4* | C4da | BDSC# 32079 <sup>72</sup> |
| *82E12-Gal4 (2<sup>nd</sup> and 3<sup>rd</sup>)* | A08n | BDSC# 40153 and this study |
| *82E12-LexA* | A08n | BDSC# 54417 |
| *NompC-LexA* | C3da, cho | BDSC# 52241 |
| *82E12-Gal4<sup>AD</sup>* | A08n split-Gal4 | <sup>24</sup> |
| *6.14.3-Gal4<sup>DBD</sup>* | A08n split-Gal4 | <sup>24</sup> |
| *LexAop-Brp<sup>short</sup>-mCherry* | Presynaptic active zone marker | <sup>32</sup> |
| *UAS-Drep2-GFP* | Postsynaptic density marker | <sup>33</sup> |
| *UAS-CD4-tdTomato* | Morphological marker | BDSC# 35841 |
| *UAS-CD4-tdGFP* | Morphological marker | BDSC# 35836 |
| *UAS-Tao<sup>RNAi</sup>* | knockdown of Tao | BDSC# 35147 |
| *UAS-Tao<sup>CA</sup>* | Expression of hyperactivate Tao | Gift from K. Harvey |
| *UAS-hTaok<sup>wt</sup>* | Expression of hTaok2 wt | This study |
| *UAS-hTaok2<sup>A135P</sup>* | Expression of hTaok2 ASD linked mutant | This study |
| *LexAop-syb-spGFP1-10, UAS-CD4-spGFP-11* | Labeling synapses with Syb-GRASP | BDSC# 64315 |
| *UAS-CsChrimson* | Optogenetic activation | BDSC# 55135 |
| *LexAop-CsChrimson* | Optogenetic activation | BDSC# 55138 |
| *UAS-GCaMP6m* | Calcium imaging | BDSC# 42748 |
| *UAS-TNTe* | Silencing neurons | BDSC#28837 |
| *LexAop-TNTe-HA* | Silencing neurons | <sup>73</sup> |
| *UAS-CaMPARI* | Ratiometric calcium integrator | BDSC# 58761 <sup>49</sup> |
| *UAS-Ras85D<sup>V12</sup>* | Expression of constitutively active Ras1 | BDSC# 4847 |
| *UAS-Rac1<sup>V12</sup>* | Expression of constitutively active Rac1 | BDSC# 6291 |

A 2^nd^ chromosome insertion of *82E12-Gal4* was generated by P-element hopping of the original insertion line (attP2 on 3^rd^ chromosome, ^28^) using Δ2,3 transposase. Human Taok2 cDNAs (wt and A135P) were cloned into pUAST-AttB carrying a C-terminal 3xflag tag. 2^nd^ chromosome transgenic lines (attP16) were generated using ϕC31-mediated transgenesis ^74^ (BestGene Inc., Chino Hills, CA, USA).

### Immunohistochemistry and confocal microscopy

Larval brains were dissected in dissection buffer (108 mM NaCl, 5 mM KCl, 4 mM NaHCO3, 1 mM NaH2PO4, 5 mM Trehalose, 10 mM Sucrose, 5 mM HEPES, 8,2 mM MgCl2, 2 mM CaCl, pH 7,4) and fixed with 4% formaldehyde/PBS for 15 min, washed with PBS and mounted on poly-L-lysine (Sigma)-coated cover slips in SlowFade Gold (ThermoFisher, Carlsbad, CA, USA). For most samples, native fluorescence was sufficiently bright and was subsequently visualized by confocal microscopy (Zeiss LSM700). Confocal z-stacks were processed in Fiji (ImageJ, NIH, Bethesda) and/or Imaris (BitPlane, Belfast, UK). For activity-dependent GFP reconstitution across synaptic partners (Syb-GRASP, Macpherson et al., 2015), larval brains were washed after dissection three times in dissection buffer containing 70 mM KCl. Larval brains were then kept in dissection buffer of 10 min to allow subsequent GFP reconstitution.

Brains were fixed in 4% formaldehyde/PBS for 15 min and native GFP fluorescence was examined by confocal microscopy. Mouse anti-Fas3 antibody was used as a sensory axon marker (7G10, 1:200, DSHB, Iowa, USA), rabbit anti-GFP (ThermoFisher, Carlsbad, CA, USA) was used to visualize presynaptic Syb-splitGFP1-10. Secondary donkey antibodies conjugated to DyLight dyes were used at 1:300 dilution (Jackson ImmunoResearch, Cambridgeshire, UK). Confocal imaging was performed as described above. All experiments were performed at least three times, examining multiple animals with consistent results.

### Synaptic marker co-localization and Syb-GRASP analysis

For co-localization analysis of pre- and postsynaptic markers (Brp^short^-mCherry and Drep2-GFP, respectively) and Syb-GRASP puncta, confocal image stacks were deconvolved using a blind deconvolution algorithm, with 10 iterations at medium noise (AutoQuant, Media Cybernetics, Rockville, MD, USA) and analyzed in Imaris (Bitplane AG, Zurich, Switzerland). Puncta were automatically detected in a region of interest using the Imaris spot function set to a size of 200 nm after background subtraction and an intensity quality threshold (Brp^short^-mCherry: 4,000, Drep2-GFP: 10,600, Syb-GRASP: 10,000) based on automatic thresholds of previous experiments with the same confocal imaging settings. Identical procedures and settings (acquisition, quality check and threshold) were used for pre- and postsynaptic puncta or Syb-GRASP for every image set as described above. Images with high noise or low contrast were discarded before analysis. Only high quality images were used for analysis to ensure consistent results, as otherwise automatic quantification was not applicable. To identify C4da-A08n neuron synapses, a spot colocalization function (MATLAB, Natick, MA, USA) was used with a distance threshold of 0.35 µm. For every animal, 4 abdominal hemisegments (a5 and a6) were analyzed and averaged. Distance threshold for synaptic contact was set to 350 nm based on estimated distances between the synaptic marker proteins similarly to previous studies ^31,34^.

### Mechanonociception assays

Mechanonociception experiments were performed with calibrated *von-Frey*-filaments (35 mN or 50 mN) and staged 3^rd^ instar larvae (96h AEL ±3h) ^23,24,75^. Larvae were carefully transferred to a 2% agar plates with a 1 ml water film and stimulated twice on mid-abdominal segments (a3-a6) within 2s. Behavioral responses (non-nociceptive, bending, rolling) were noted and only behavioral responses to the 2^nd^ stimulus were analyzed and plotted. Staging and experiments were done in a blinded fashion and randomized fashion.

### Optogenetic behavioral assays

For optogenetic behavior, larvae were grown in darkness on grape agar plates with yeast paste containing 5 mM all-*trans*-retinal. Staged 3^rd^ instar larvae (96h ±3h AEL) were carefully transferred under low red light conditions to a 2% agar plates with a 1 ml water film. CsChrimson was activated with 625 nm light (760 µW/cm²) for 5 s. Videos were taken during the experiment and analyzed using the Fiji cell counter plugin (ImageJ, NIH, Bethesda). Staging, experiments and analyses were done in a blinded fashion. Rolling was defined as at least one complete 360° roll. Bending was defined as a c-shape like twitching, typically seen before rolling behavior, and not to be confused with other described bending behavior (Jovanic et al. 2016). Turning behavior describes head turning and thereby a direction changes of locomotion. All behavioral assays and analyses were performed in a blinded and randomized fashion.

### Cold Plate Assay

For the cold plate assay, 20-30 larvae (96h AEL ±3h) were placed on a 7.2 mm × 11.2 mm × 1 mm brass metal plate, which was coated with a 1 mm 2% agar and 1 ml water film. The plate was put on the heating/cooling block of a PCR Thermo-Cycler (Biometra TGradient, Biomedizinische Analytik GmbH, Göttingen, Germany). A program was established so the temperature on the agar fell from 25 °C to 3 °C within 50 s after staying at 25 °C for 15 s (15 s at 25 °C, cooling to −3 °C). Temperature of the agar surface was monitored and video-captured during the whole experiment (GTH 1170 digital thermometer, Greisinger electronic, Remscheid, Germany). Videos were taken using a digital camera (Basler ace acA2040-25gm, Basler AG, Ahrensburg, Germany) and FrameGrabber software (StreamPix, NorPix Inc., Montreal, Canada) and analyzed using the Fiji cell counter plugin (ImageJ, NIH, Bethesda). Experiments and analyses were done in a blinded fashion. Stop and turning behavior describes stopping of typical larval locomotion with subsequent head turning behavior. Contraction behavior was defined by contraction of the larval body in an unbent, straight fashion. Contraction and bending was defined by contraction behavior with simultaneous c-shape like bending. Contraction and rolling was defined by contraction behavior with at least one simultaneous 360° roll. Rolling was defined as at least one complete 360° roll. Bending was defined as c-shaped twitching, not to be confused with other described bending behavior (Jovanic et al. 2016). Response categories were defined and numbered according to progressively stronger behavioral responses (1 = crawling, 2 = stop & turn, 3 = contraction, 4 = contraction & bending, 5 = contraction & rolling, 6 = bending, 7 = rolling). The highest response category of an individual animal was defined as the observed behavior corresponding to the highest numerical value defined above to describe changes from C3da to C4da neuron-dependent responses. All behavioral assays and analyses were performed in a blinded and randomized fashion.

### GCaMP6m calcium imaging

Staged 3^rd^ instar larvae (96h (+-3) AEL) were partially dissected in physiological saline buffer (120 mM NaCl, 3 mM KCl, 10 mM Trehalose, 10 mM Glucose, 10 mM Sucrose, 10 mM NaHCO3, 4 mM MgCl2, 1,5 mM CaCl, 10 mM HEPES, pH 7,25) and pinned on a Sylgard plate to expose the VNC. A08n neuron somata expressing Gcamp6m were live imaged by confocal microscopy with a 40x/NA1.0 water objective (Zeiss LSM700, Zeiss, Oberkochen, Germany). Activation of sensory neurons induced by C3da or C4da-specific CsChrimson activation was achieved using a 635 nm LED (Mightex, Pleasanton, CA, USA) filament with maximum output of 70 µW/cm². Confocal time series were taken at 4.1 frames/s (320×320 pixels). A08n somata were focused and after 20 frames of stable imaging, the 635 nm LED was activated for 5 s. Times series files were analyzed in Fiji/ImageJ using image registration (StackReg plugin) to correct for VNC movement and subsequent quantification of GCaMP6m signal intensity in the soma using the Time Series Analyzer V3 plugin (ImageJ). Baseline (F0) was determined by the average of 15 frames before activation. Relative maximum intensity change (ΔFmax) of Gcamp6m fluorescence was calculated after normalization to baseline.

### CaMPARI calcium integrator assay

CaMPARI, a photoconvertible calcium integrator ^49^, was converted with UV light to measure A08n neuronal activity in the presence of a 4°C cold stimulus. The ratio of photoconversion correlates with calcium levels in neurons during the time window defined by the UV conversion light. 96h AEL old larvae were put on a 6 cm grape agar petri dish. A drop of 80 µl cold water at 4 °C was applied and the larvae were exposed to 20s of photo-conversion light (385 nm, 0.537 mW/mm²). Larval brains were dissected, fixed in 4% Formaledhyde/PBS solution for 15min, and imaged with a confocal microscope. For quantification of the conversion ratio, maximum intensity projections of the acquired z-stacks were analyzed (A08n soma region, equal stack size). Intensities of the red and green fluorescent CaMPARI forms were measured in A08n somata (ImageJ, NIH, Bethesda) to acquire Fred/Fgreen ratios.

### Electron microscopy analysis of C4da-A08n synapses

EM samples were processed and imaged as described previously (Hu et al. 2017). Drep2-GFP and Brp^short^-mCherry were expressed in A08n and C4da neurons to specifically visualize C4da presynaptic active zones and A08n postsynaptic densities, respectively (27H06-LexA>LexAop-Brp^short^-mCherry; 82E12-Gal4>UAS-Drep2-GFP). Larvae (96h AEL) were dissected in 0.1 M Sorensen’s Phosphate Buffer (SPB, Electron Microscopy Sciences, Hatfield, PA, USA), fixed with periodate-lysine-paraformaldehyde (PLP, Electron Microscopy Sciences, Hatfield, PA, USA) for 20 min followed by fixation in 0.01% glutaraldehyde in SPB for another 30 min at RT. Free aldehyde was quenched with 1 mM glycine in SPB for 10 min. To ensure antibody penetration of the blood-brain barrier, it was necessary to carefully puncture the brain’s surface with an electrolytically sharpened tungsten needle. Subsequently, 0.3% H2O2 was applied for 30 min to block endogenous peroxidase activity. After washing and blocking in a buffer (SPB, 0,25% Saponin (Sigma-Aldrich, St. Louis, MO, USA)) brains were stained with rabbit αGFP (1:500, #A-11122, ThermoFisher, Waltham, MA, USA) and rabbit αdsRed (1:500, #632496, ClonTech, Mountain View, CA, USA) in blocking solution, followed by the VectaStain ABC kit procedure with a biotinylated anti-rabbit secondary antibody and HRP (Vector Laboratories, Peterborough, UK). Samples were incubated in SPB containing 0.5 mg/mL DAB and 0,003% H2O2 to visualize HRP activity. Stained specimens were postfixed in 2% GA in SPB for 1h and then treated with 1% OsO4 for 1 h on ice. After an ethanol dehydration series, the specimens were embedded in epon (Roth, Karlsruhe, Germany). We cut 0.5 µm semi-thin cross-sections of the VNC and examined them under a light microscope to find DAB-stained areas with labeled neuronal structures. Ultrathin sections of 60 nm were examined and imaged on an EM902 (Zeiss, Oberkochen, Germany). Serial sections from multiple animals were examined and images were taken with a TRS 2K digital camera (A. Tröndle, Moorenweis, Germany). Multiple section series were then analyzed for C4da active zones and A08n postsynaptic compartments. Individual labeled compartments from multiple animals (n=4) were counted and the number of direct C4da-A08n synaptic contacts was determined and given as a ratio to the total of identified C4da pre- or A08n postsynapses.

### Quantification and statistical analysis

Statistical analysis was done with Prism 6 (Graphpad, San Diego, CA, USA). All shown whiskers represent standard deviation. Appropriate statistical tests were applied and are mentioned in figure legends. Sample sizes were chosen to be similar to be similar to previous publications (Mosca et al., 2014;Ohyama et al., 2015; Burgos et al. 2018; Hu et al., 2017). For normal distributed data, unpaired two-tailed Student’s *t*-test (two groups), One Way ANOVA (multiple comparison) and χ² tests (two groups with more than two categorized behavioral data, Fisher’s exact test for two categorized behavioral data) were used to compare groups. For multiple comparisons, Dunnett’s *post-hoc* test was performed to compare several groups with a control group, Sidak’s *post-hoc* test was performed to compare preselected pairs.

## Acknowledgements

The authors thank G. Tavosanis for LexAop-Brpshort-mCherry, K. Harvey for UAS-TaoCA, C.H. Lee and R. Benton for LexAop-TNTe lines; J.Z. Parrish for comments on the manuscript, J. Felix Evers for discussion of unpublished results. Stocks obtained from the Bloomington Drosophila Stock Center (NIH P40OD018537) were used in this study. This work was supported by ERANET-NEURON (BMBF 01EW1910, to P.S. and F.C.d.A.), JPND (BMBF 01ED1806 to F.C.d.A.), the Landesforschungsförderung Hamburg (LFF-LV17/A2, to P.S. and F.C.d.A.) and the Deutsche Forschungsgemeinschaft (DFG grants SO1337/2-1 and SO1337/4-1 to P.S., CA1495/4-1 to F.C.d.A.).

## Author contributions

F.M.T and P.S designed experiments, F.M.T, M.S.G, L.P, M.M.P, C.H. and B.S. collected confocal imaging data. F.M.T collected Ca2+ imaging and behavioral data. D.W., E.S. and M.S collected ultrastructural data. M.R., M.P, S.S. and F.C.d.A. provided essential reagents. All analyses and statistics were performed by F.M.T. F.M.T and P.S wrote the manuscript.

## Supplementary Information

**Supplementary Figure 1.**
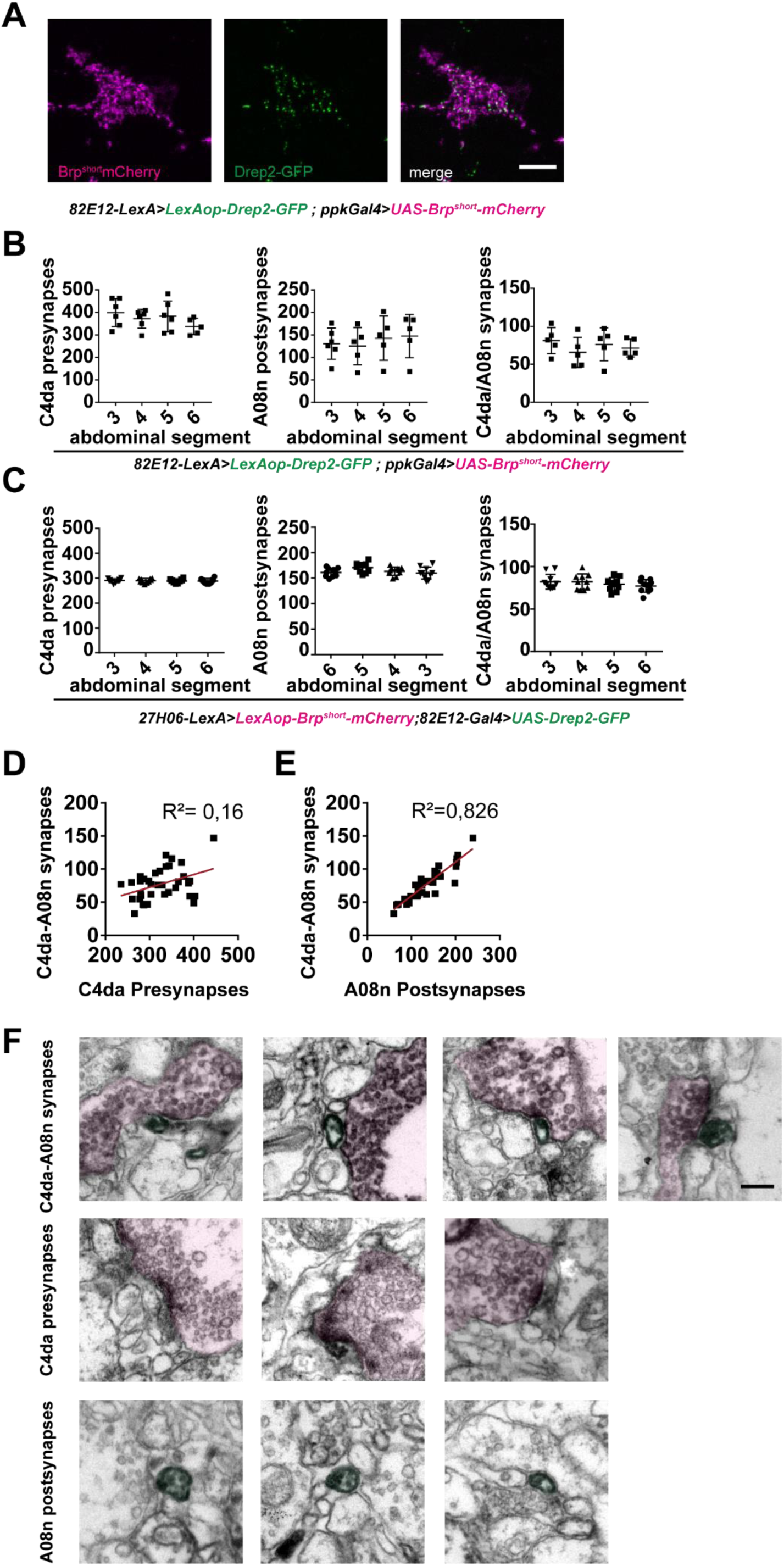
(A) Larval VNC hemisegment (96h AEL) with expression of Brp^short^-mCherry in C4da (magenta), Drep2-GFP in A08n (green) and merge (*82E12-LexA>LexAop-Drep2-GFP; ppk-Gal4>UAS-Brp^short^-mCherry*). Scale bar = 5 µm. (B) Quantification of C4da presynapses, A08n postsynapses and C4da-A08n synpases in abdominal segments 3-6 using *82E12-LexA>LexAop-Drep2-GFP; ppk-Gal4>UAS-Brp^short^-mCherry*. n = 6. (C) Quantification of C4da presynapses, A08n postsynapses and C4da-A08n synpases in abdominal segments 3-6 using *27H06-LexA>LexAop-Brp^short^-mCherry; 82E12-Gal4>UAS-Drep2-GFP*. n = 10. (D) Correlation between C4da-A08n synapses and C4da presynapses with linear regression. R²=0.16, n = 40 hemisegments from 10 animals (E) Correlation between C4da-A08n synapses (colocalization) plotted against A08n postsynapses with linear regression. R²=0.826, n = 40 hemisegments from 10 animals. (F) EM images showing DAB-labelled C4da presynapses (colored in magenta) connected to A08n postsynaptic compartments (colored in green), and C4da or A08n synaptic compartments without labelled counterparts. Scale bar = 200 nm.

**Supplementary Figure 2.**
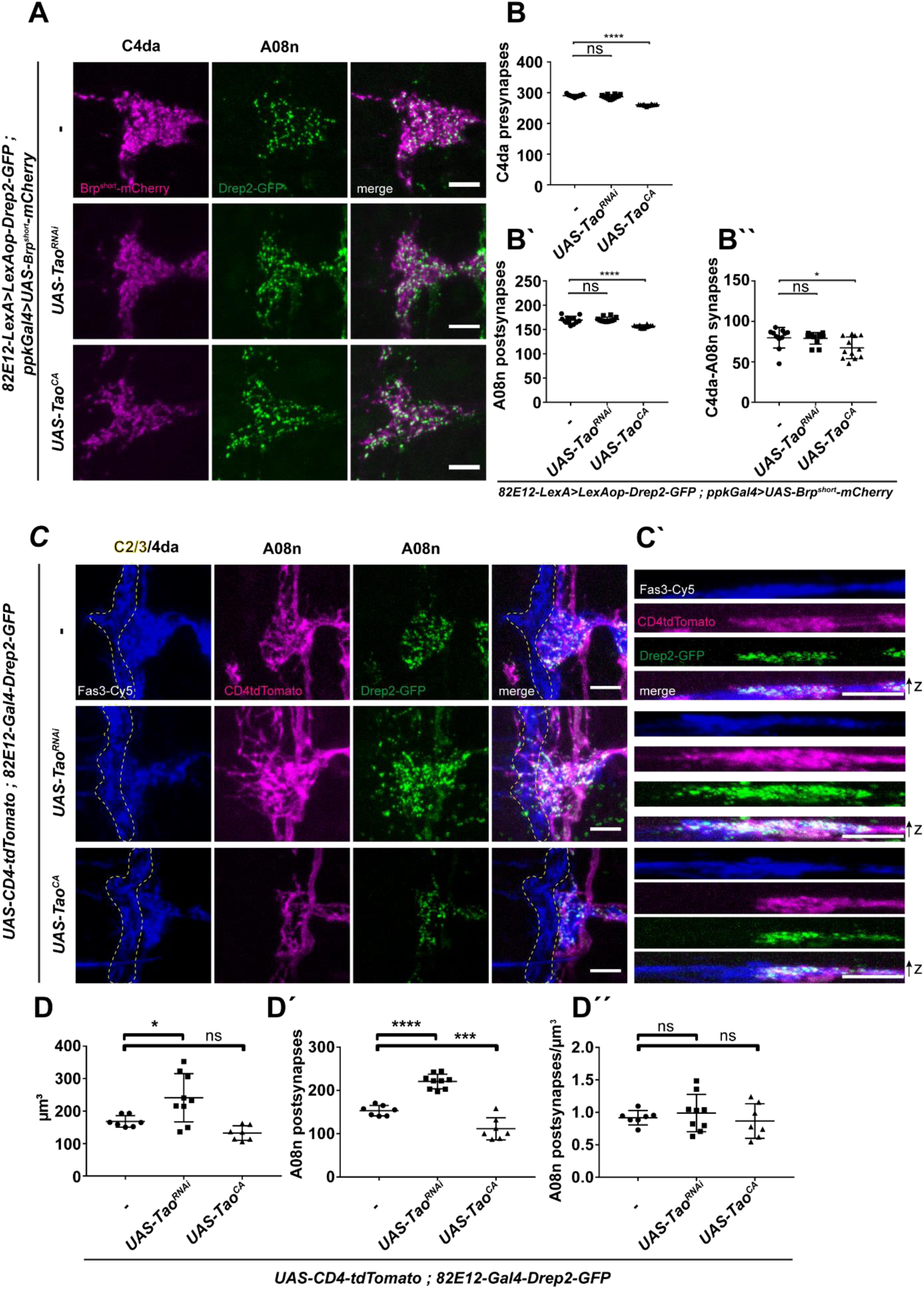
(A) Confocal images of larval VNC hemisegments (96h AEL) in *control*, or with *Tao^RNAi^* and Tao^CA^ expression in C4da neurons using synaptic markers labeling C4da presynapses (magenta) and A08n postsynapses (green). Scalebar = 5 µm. (B) Quantification of C4da pre-, (B’) A08n post- and (B’’) colocalized C4da-A08n synaptic markers in control or with *Tao^RNAi^* and Tao^CA^ expression in C4da neurons. \**P<*0.05, **** *P* < 0.0001. ±SD, ANOVA with multiple comparisons and Dunnett’s *post-hoc* test (for exact *P* values and statistics see Supplemental Table 1). Control n = 11, *UAS-Tao^RNAi^* n = 12, *UAS-Tao^CA^* n = 12. (C) Confocal images of hemisegments in control or with *Tao^RNAi^* and Tao^CA^ expression in A08n neurons. A08n neurons were labeled using the morphological marker *CD4-tdTomato* (magenta) and postsynaptic marker *Drep2-GFP* (green), together with anti-Fas3 immunostaining labeling C2da, C3da and C4da sensory axons (blue). C2da/C3da domain is indicated by yellow dotted line. Scale bar = 5 µm. (C’) XZ projections of each channel in (C) are shown. Scale bar = 5 µm. (D) Quantification of dendrite volume, (D’) A08n postsynapses and (D’’) A08n postsynapse/volume ratio per hemisegment in control or with *Tao^RNAi^* and Tao^CA^ expression in A08n neurons. \**P<*0.05, \*\**P<*0.005, **** *P* < 0.0001. ±SD, ANOVA with multiple comparisons and Dunnett’s *post-hoc* test (for exact *P* values and statistics see Supplemental Table 1). Control n = 7, *UAS-Tao^RNAi^* n = 9, *UAS-Tao^CA^* n = 7.

**Supplementary Figure 3.**
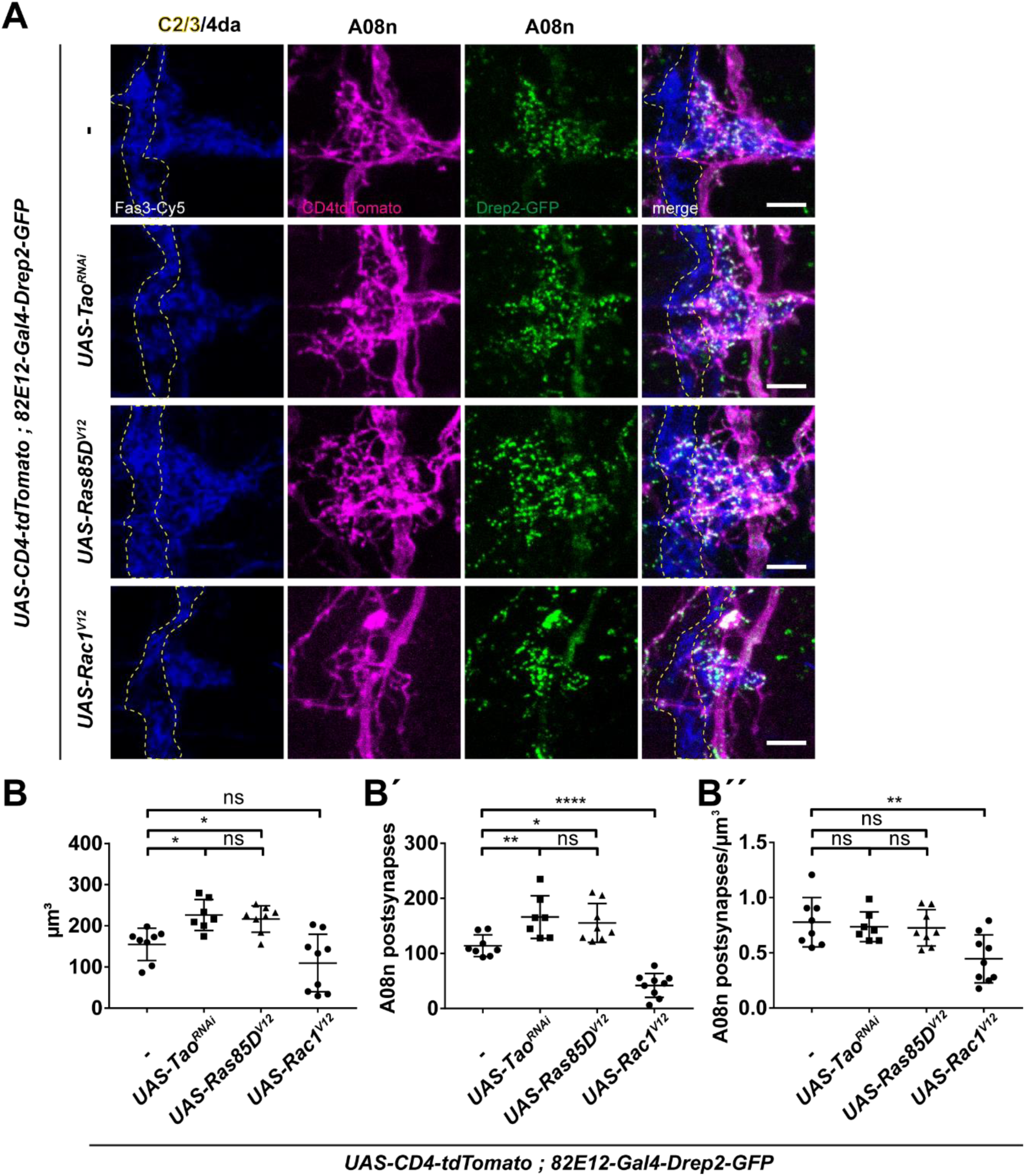
Ras1 and Rac1 overexpression differentially regulate dendrite growth and synaptogenesis of A08n neurons. (A) Confocal images of larval VNC hemisegments (96h AEL) in control or with Tao^RNAi^, Ras85D^V12^ and Rac1^V12^ expression in A08n neurons. A08n neurons were labeled using the morphological marker CD4-tdTomato (magenta) and postsynaptic marker Drep2-GFP (green), together with anti-Fas3 immunostaining labeling C2da, C3da and C4da sensory axons (blue). C2da/C3da domain is indicated by yellow dotted line. Scale bar = 5 µm. (B) Quantification of dendrite volume, (B’) A08n postsynapses and (B’’) postsynapse/volume ratio per hemisegment in control or with Tao^RNAi^, Ras85D^V12^ and Rac1^V12^ expression in A08n neurons. * P<0.05, ** P<0.01, **** P < 0.0001. ±SD, ANOVA with multiple comparisons and Dunnett’s post-hoc test (for exact P values and statistics see Supplemental Table 1). control n = 8, UAS-Tao^RNAi^ n = 7, UAS-Ras85D^V12^ n = 8, UAS-Rac1^V12^ n = 9.

**Supplementary Figure 4.**
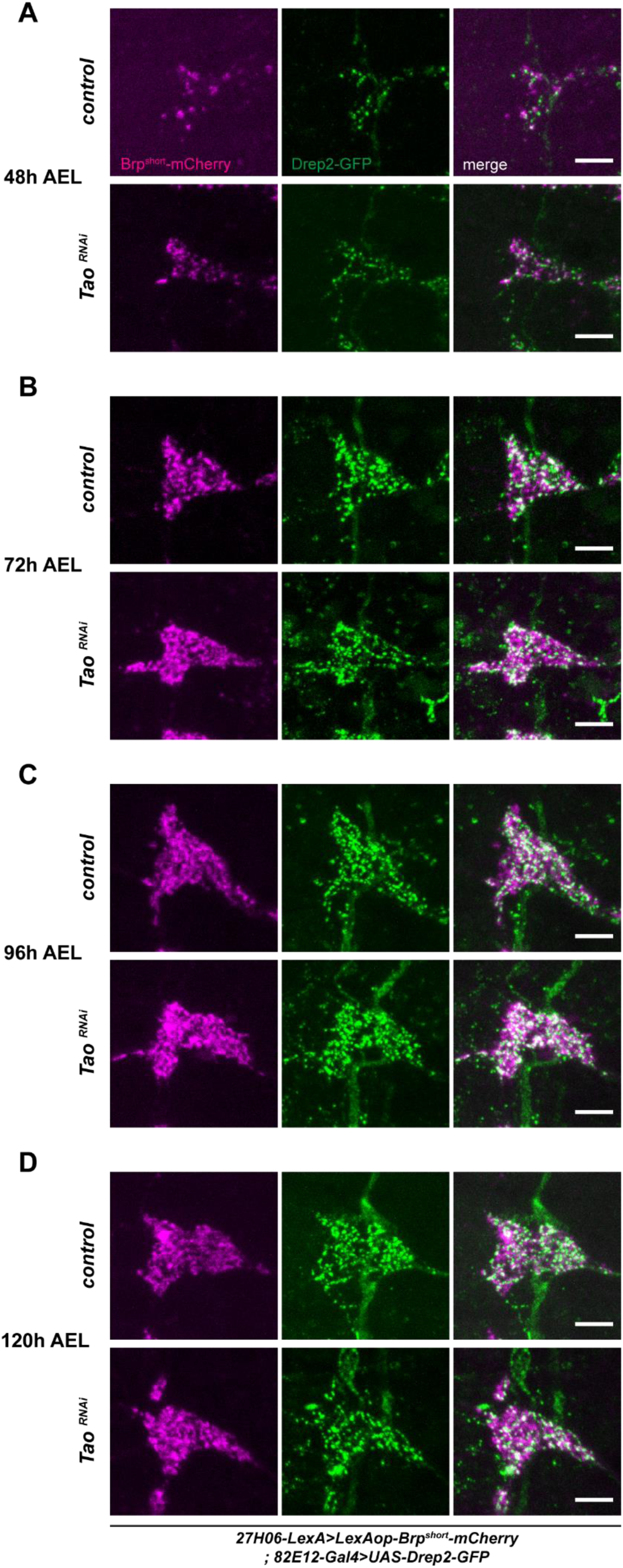
Confocal images of representative larval hemisegments during development from 48 AEL until 120h AEL in control or with *Tao^RNAi^* expression in A08n neurons. Synaptic markers labeling C4da presynapses (magenta), A08n postsynapses (green) are shown at (A) 48h, (B) 72h, (C) 96h and (D) 120h AEL. Scale bar = 5 µm.

**Supplementary Figure 5.**
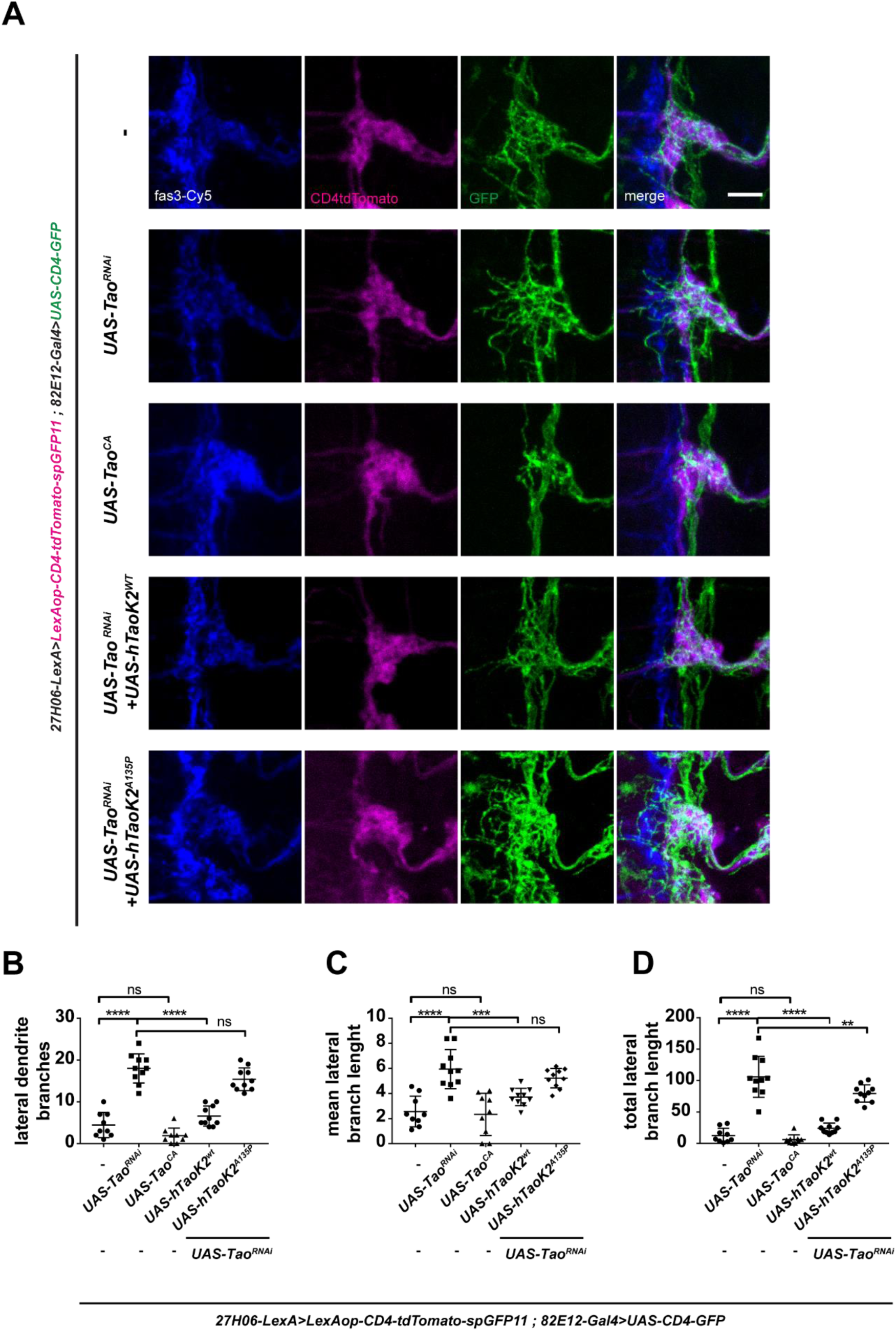
(A) Confocal Images of larval VNC hemisegments (96h AEL) in control, with *Tao^RNAi^* and Tao^CA^, or co-expression of *Tao^RNAi^* with hTaoK2^wt^ or hTaoK2^A135P^ in A08n neurons. Images show anti-Fas3 immunostaining labeling C2da, C3da and C4da sensory axons (blue), with anatomical marker expression to label C4da axons (CD4-tdTomato, magenta) and A08n neurons (CD4-tdGFP, green). Scale bar = 5 µm. (B-D) Quantification of A08n neuron ectopic dendritic branches per hemisegment laterally displaced exiting the C4da domain (lateral dendrite branches). (B) Lateral dendrite numbers, (C) mean branch length and (D) total lateral branch length in control, with *Tao^RNAi^* and Tao^CA^, or co-expression of *Tao^RNAi^* with hTaoK2^wt^ or hTaoK2^A135P^ in A08n neurons. Control: n = 9, *UAS-Tao^RNAi^*: n = 10, *UAS-Tao ^CA^*: n = 9, *UAS-Tao^RNAi^ + UAS-hTaoK2 ^wt^*: n = 10, *UAS-Tao^RNAi^ + UAS-hTaoK2 ^A135P^* n = 10. \*\*\**P<*0.001, **** *P* < 0.0001. ±SD, ANOVA with multiple comparisons and Dunnett’s *post-hoc* test (for exact *P* values and statistics see Supplemental Table 1).

**Supplementary Figure 6.**
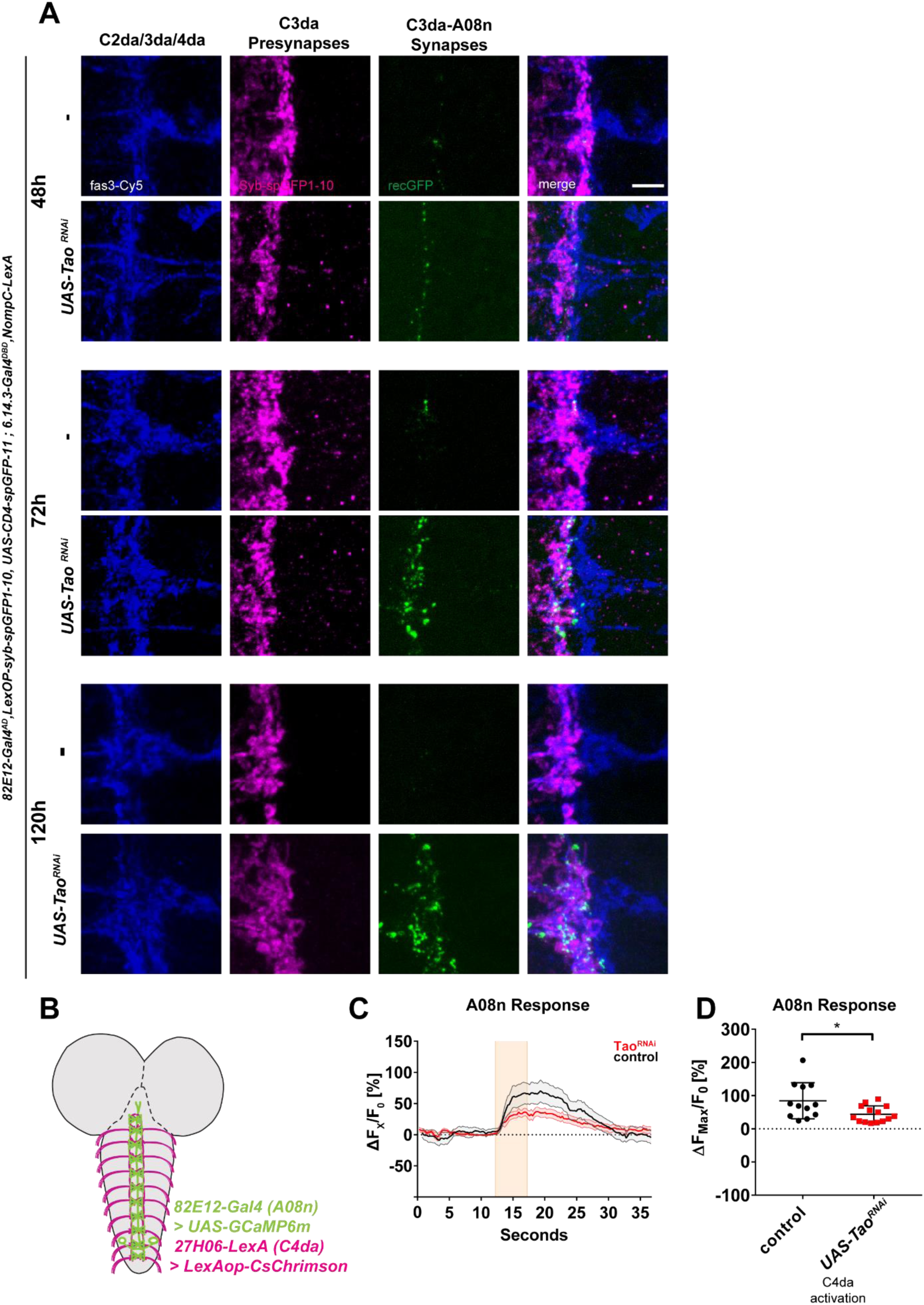
(A) Confocal images of Syb-GRASP-labeled C3da-A08n synapses (48, 72 and 120h AEL). Representative images of larval VNC hemisegments in control or with *Tao^RNAi^* expression in A08n neurons showing anti-Fas3 labeling of C2da, C3da and C4da sensory axons (blue), presynaptic Syb-spGFP1-10 expressed in C3da (magenta) and reconstituted native GFP signal marking C3da-A08n Synapses (green). Scale bar = 5 µm. (B) Schematic larval brain showing A08n neurons (green) and C4da sensory dendrite VNC projections (magenta) and indicating expression of *UAS-GCamp6m* in A08n and *LexAop-CsCrimson* in C4da neurons. (C) Calcium responses of Gcamp6m-expressing A08n neurons after optogenetic activation of C4da neurons using CsChrimson (5s, 630 nm, indicated by shaded area), with or without *Tao^RNAi^* expression in A08n neurons. Data show mean change in percent [(ΔF/F0)-1, (±SEM indicated by shaded regions]. *Control* n = 11, *UAS-Tao^RNAi^* n = 9. (D) Quantification of maximum A08n responses to C4da activation in percent [(ΔFMax/F0)-1)] comparing control and *Tao^RNAi^* expression in A08n neurons. Control n=11, *UAS-Tao^RNAi^* n=9. *P*= 0,0199. ±SD. Unpaired two-tailed *t*-test.

**Supplementary Figure 7.**
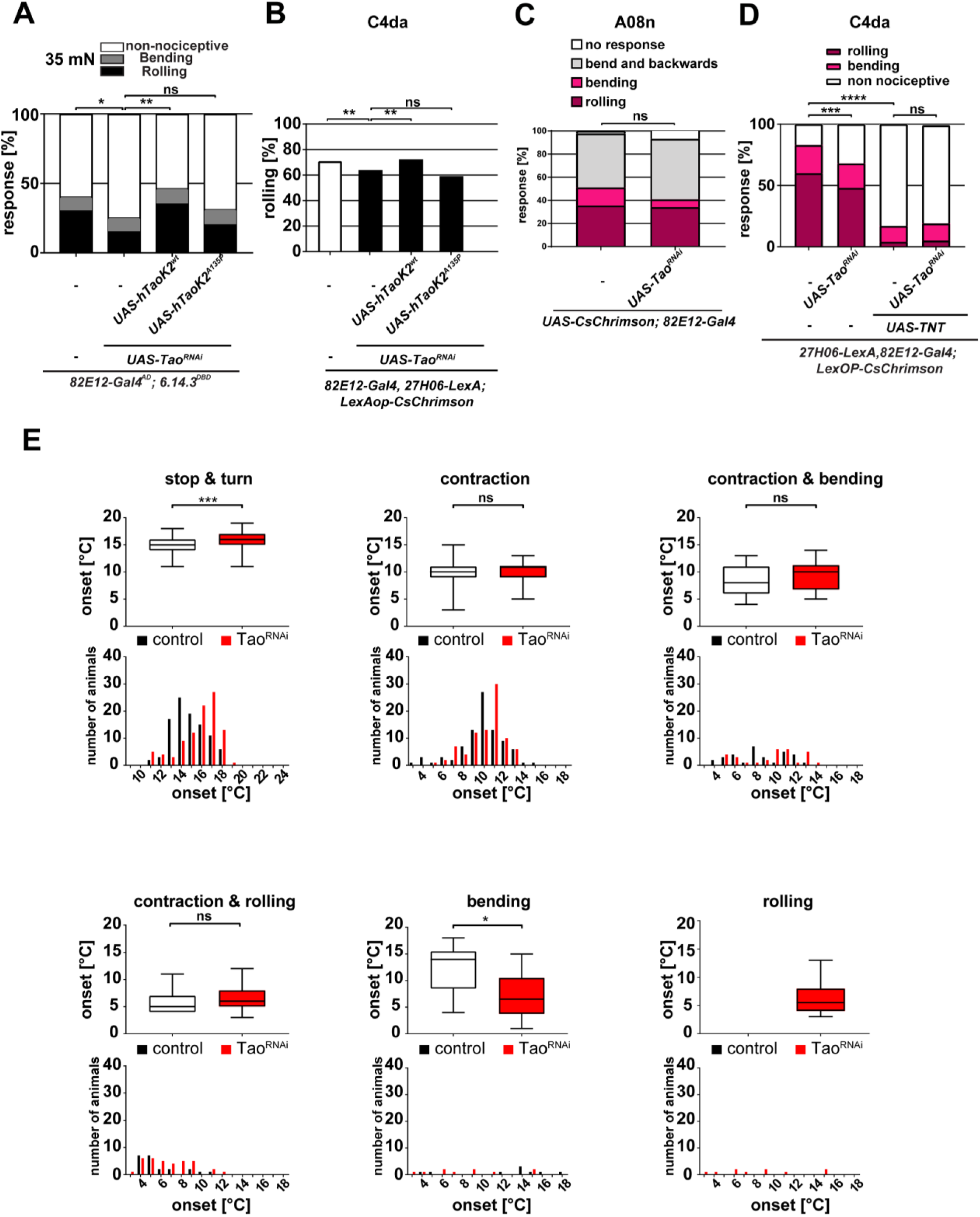
(A) Mechanonociceptive behavioral response of third instar larvae (96h AEL) in control or with *Tao^RNAi^* without or with co-expression of hTaoK2^wt^ or hTaoK2^A135P^ in A08n neurons. Responses to the second mechanical stimulation with a 35 mN *von Frey* filament are shown (Nociceptive rolling and bending or non-nociceptive responses). \**P* < 0.05, \*\*\**P* < 0.001. *Control* n = 100, *UAS-Tao^RNAi^* n = 98, *UAS-Tao^RNAi^ + UAS-hTaoK2^wt^* n = 99, *UAS-Tao^RNAi^ + UAS-hTaoK2^A135P^* n = 99. Control vs *UAS-Tao^RNAi^*: *P =* 0.0439, *UAS-Tao^RNAi^* vs *UAS-Tao^RNAi^ + UAS-hTaok2*^wt^: *P =* 0.0037, *UAS-Tao^RNAi^* vs *UAS-Tao^RNAi^ + UAS-hTaok2^A135P^*: *P =* 0.6268, χ^2^ test.(B) Behavioral responses of third instar larvae (96h AEL) to optogenetic activation of C4da neurons expressing *CsChrimson* (5 s, 625 nm light pulse) in control, or *Tao^RNAi^* without or with co-expression of hTaoK2^wt^ or hTaoK2^A135P^ in A08n neurons. \*\**P* < 0.005. *Control*: n = 95, *UAS-Tao^RNAi^*: n = 91, *UAS-Tao^RNAi^ + UAS-hTaoK2^wt^*: n = 80, *UAS-Tao^RNAi^ + UAS-hTaoK2^A135P^*: n = 98. *Control vs UAS-Tao^RNAi^*: *P =* 0,0049, *UAS-Tao^RNAi^* vs *UAS-Tao^RNAi^ + UAS-hTaok2*^wt^: *P =* 0.0016, *UAS-Tao^RNAi^* vs *UAS-Tao^RNAi^ + UAS-hTaok2^A135P^*: *P =* 0.2667, χ^2^ test (C) Behavioral responses of third instar larvae (96h AEL) to optogenetic activation of A08n neurons expressing CsChrimson (5 s, 625 nm light pulse) in control or *Tao^RNAi^* expression in A08n neurons. *Control*: n = 95, *UAS-Tao^RNAi^*: n = 91 *P =* 0.1005, χ^2^ test. (D) Behavioral responses of third instar larvae (96h AEL) to optogenetic activation of C4da neurons expressing CsChrimson (5 s, 625 nm light pulse) in control, or with *Tao^RNAi^* and TNT expression in A08n neurons. \*\**P* < 0.005, \*\*\*\**P* < 0.0001. *Control* n = 206, *UAS-Tao^RNAi^* n = 185, *UAS-TNTe* n = 288, *UAS-Tao^RNAi^ + UAS-TNTe* n = 102. *Control* vs *UAS-Tao^RNAi^*: *P =* 0.0005, *Control* vs *UAS-TNTe P =* >0.0001, *UAS-TNTe* vs *UAS-Tao^RNAi^ + UAS-TNTe P =* 0.7307, χ^2^ test. (E) Temperature-dependent onset and animal distribution of the observed behaviors: stop & turn, contraction, contraction & bending, contraction & rolling, bending and rolling. Stop & turn: control n = 98, *UAS-Tao^RNAi^* n = 96. Contract: *control* n = 87, *UAS-Tao^RNAi^* n = 84. Contract & bending *control* n = 31, *UAS-Tao^RNAi^* n = 30. Contraction & rolling: *control* n = 22, *UAS-Tao^RNAi^* n = 35. Bending: *control* n = 9, *UAS-Tao^RNAi^* n = 11. Rolling: *control* n = 0, *UAS-Tao^RNAi^* n = 20 *\*P =* >0.05 \**\*\*P =* >0.001 \**\*\*\*P =* >0.0001. ±SD Unpaired two-tailed *t*-test. Box and whisker plots depict median, 25^th^ and 75^th^ percentile (lower and upper end of box, respectively), and 5^th^ and 95^th^ percentile (lower and upper whiskers, respectively) of the data points.

## References

1. Tau, G. Z. & Peterson, B. S. Normal development of brain circuits. Neuropsychopharmacology 35, 147–168 (2010).

2. Davis, G. W. Homeostatic signaling and the stabilization of neural function. Neuron 80, 718–728 (2013).

3. Turrigiano, G. Too Many Cooks? Intrinsic and Synaptic Homeostatic Mechanisms in Cortical Circuit Refinement. Annu. Rev. Neurosci. 34, 89–103 (2011).

4. Riccomagno, M. M. & Kolodkin, A. L. Sculpting Neural Circuits by Axon and Dendrite Pruning. Annu. Rev. Cell Dev. Biol. 31, 779–805 (2015).

5. Bleckert, A. & Wong, R. O. L. Identifying roles for neurotransmission in circuit assembly: insights gained from multiple model systems and experimental approaches. Bioessays 33, 61–72 (2011).

6. Gerhard, S., Andrade, I., Fetter, R. D., Cardona, A. & Schneider-Mizell, C. M. Conserved neural circuit structure across Drosophila larval development revealed by comparative connectomics. Elife 6, 1–17 (2017).

7. Keshishian, H. et al. Cellular mechanisms governing synaptic development inDrosophila melanogaster. J. Neurobiol. 24, 757–787 (1993).

8. Zwart, M. F., Randlett, O., Evers, J. F. & Landgraf, M. Dendritic growth gated by a steroid hormone receptor underlies increases in activity in the developing Drosophila locomotor system. Proc. Natl. Acad. Sci. U. S. A. 110, E3878–87 (2013).

9. Grueber, W. B., Jan, L. Y. & Jan, Y. N. Tiling of the Drosophila epidermis by multidendritic sensory neurons. Development 129, 2867–78 (2002).

10. Parrish, J. Z., Xu, P., Kim, C. C., Jan, L. Y. & Jan, Y. N. The microRNA bantam Functions in Epithelial Cells to Regulate Scaling Growth of Dendrite Arbors in Drosophila Sensory Neurons. Neuron 63, 788–802 (2009).

11. Guan, B., Hartmann, B., Kho, Y.-H., Gorczyca, M. & Budnik, V. The Drosophila tumor suppressor gene, dlg, is involved in structural plasticity at a glutamatergic synapse. Curr. Biol. 6, 695–706 (1996).

12. Davis, G. W. & Goodman, C. S. Synapse-specific control of synaptic efficacy at the terminals of a single neuron. Nature 392, 82–86 (1998).

13. Choi, B. J. et al. Miniature Neurotransmission Regulates Drosophila Synaptic Structural Maturation. Neuron 82, 618–634 (2014).

14. Rasse, T. M. et al. Glutamate receptor dynamics organizing synapse formation in vivo. Nat. Neurosci. 8, 898–905 (2005).

15. Eichler, K. et al. The complete connectome of a learning and memory centre in an insect brain. Nature 548, 175–182 (2017).

16. Miroschnikow, A. et al. Convergence of monosynaptic and polysynaptic sensory paths onto common motor outputs in a Drosophila feeding connectome. Elife 7, 1–23 (2018).

17. Schlegel, P. et al. Synaptic transmission parallels neuromodulation in a central food-intake circuit. Elife 5, 462–465 (2016).

18. Ohyama, T. et al. A multilevel multimodal circuit enhances action selection in Drosophila. Nature 520, 633–639 (2015).

19. White, J. G., Southgate, E., Thomson, J. N. & Brenner, S. The Structure of the Nervous System of the Nematode Caenorhabditis elegans. Philos. Trans. R. Soc. B Biol. Sci. 314, 1–340 (1986).

20. Tracey, W. D. et al. painless, a Drosophila gene essential for nociception. Cell 113, 261–73 (2003).

21. Hwang, R. Y. et al. Nociceptive neurons protect Drosophila larvae from parasitoid wasps. Curr Biol 17, 2105–2116 (2007).

22. Zhong, L. et al. Thermosensory and nonthermosensory isoforms of Drosophila melanogaster TRPA1 reveal heat-sensor domains of a thermoTRP Channel. Cell Rep. 1, 43–55 (2012).

23. Almeida-Carvalho, M. J. et al. The Ol 1 mpiad: concordance of behavioural faculties of stage 1 and stage 3 Drosophila larvae. J. Exp. Biol. 220, 2452–2475 (2017).

24. Hu, C. et al. Sensory integration and neuromodulatory feedback facilitate Drosophila mechanonociceptive behavior. Nat. Neurosci. 20, 1085–1095 (2017).

25. Burgos, A. et al. Nociceptive interneurons control modular motor pathways to promote escape behavior in Drosophila. Elife 7, e26016 (2018).

26. Yoshino, J., Morikawa, R. K., Hasegawa, E. & Emoto, K. Neural Circuitry that Evokes Escape Behavior upon Activation of Nociceptive Sensory Neurons in Drosophila Larvae. Curr. Biol. 1–6 (2017). doi:10.1016/j.cub.2017.06.068

27. Kaneko, T. et al. Serotonergic Modulation Enables Pathway-Specific Plasticity in a Developing Sensory Circuit in Article Serotonergic Modulation Enables Pathway-Specific Plasticity in a Developing Sensory Circuit in Drosophila. Neuron 95, 623–638.e4 (2017).

28. Vogelstein, J. T. et al. Discovery of brainwide neural-behavioral maps via multiscale unsupervised structure learning. Science 344, 386–92 (2014).

29. Takagi, S. et al. Divergent Connectivity of Homologous Command-like Neurons Mediates Segment-Specific Touch Responses in Drosophila. Neuron 96, 1373–1387.e6 (2017).

30. Macpherson, L. J. et al. Dynamic labelling of neural connections in multiple colours by trans-synaptic fluorescence complementation. Nat. Commun. 6, 10024 (2015).

31. Mosca, T. J. & Luo, L. Synaptic organization of the Drosophila antennal lobe and its regulation by the Teneurins. Elife 3, 1–29 (2014).

32. Leiss, F. et al. Characterization of dendritic spines in the Drosophila central nervous system. Dev. Neurobiol. 69, 221–34 (2009).

33. Andlauer, T. F. M. et al. Drep-2 is a novel synaptic protein important for learning and memory. Elife 3, 1–24 (2014).

34. Sheng, C. et al. Experience-dependent structural plasticity targets dynamic filopodia in regulating dendrite maturation and synaptogenesis. Nat. Commun. 9, 3362 (2018).

35. Schuster, C. M., Davis, G. W., Fetter, R. D. & Goodman, C. S. Genetic dissection of structural and functional components of synaptic plasticity. II. Fasciclin II controls presynaptic structural plasticity. Neuron 17, 655–667 (1996).

36. Liu, T., Rohn, J. L., Picone, R., Kunda, P. & Baum, B. Tao-1 is a negative regulator of microtubule plus-end growth. J. Cell Sci. 123, 2708–2716 (2010).

37. Ball, R. W. et al. Retrograde BMP Signaling Controls Synaptic Growth at the NMJ by Regulating Trio Expression in Motor Neurons. Neuron 66, 536–549 (2010).

38. Koh, Y.-H., Ruiz-Canada, C., Gorczyca, M. & Budnik, V. The Ras1-mitogen-activated protein kinase signal transduction pathway regulates synaptic plasticity through fasciclin II-mediated cell adhesion. J. Neurosci. 22, 2496–504 (2002).

39. Richter, M. et al. Altered TAOK2 activity causes autism-related neurodevelopmental and cognitive abnormalities through RhoA signaling. Mol. Psychiatry (2018). doi:10.1038/s41380-018-0025-5

40. Yadav, S. et al. TAOK2 Kinase Mediates PSD95 Stability and Dendritic Spine Maturation through Septin7 Phosphorylation. Neuron 93, 379–393 (2017).

41. de Anda, F. C. et al. Autism spectrum disorder susceptibility gene TAOK2 affects basal dendrite formation in the neocortex. Nat. Neurosci. 15, 1022–31 (2012).

42. Grueber, W. B. et al. Projections of Drosophila multidendritic neurons in the central nervous system: links with peripheral dendrite morphology. Development 134, 55–64 (2007).

43. Klapoetke, N. C. et al. Independent optical excitation of distinct neural populations. Nat. Methods 11, 338–46 (2014).

44. Tsubouchi, A., Caldwell, J. C. & Tracey, W. D. Dendritic filopodia, Ripped Pocket, NOMPC, and NMDARs contribute to the sense of touch in Drosophila larvae. Curr. Biol. 22, 2124–34 (2012).

45. Turner, H. N. et al. The TRP Channels Pkd2, NompC, and Trpm Act in Cold-Sensing Neurons to Mediate Unique Aversive Behaviors to Noxious Cold in Drosophila. Curr. Biol. 26, 3116–3128 (2016).

46. Yan, Z. et al. Drosophila NOMPC is a mechanotransduction channel subunit for gentle-touch sensation. Nature 493, 221–5 (2013).

47. Turner, H. N., Patel, A. A., Cox, D. N. & Galko, M. J. Injury-induced cold sensitization in Drosophila larvae involves behavioral shifts that require the TRP channel Brv1. PLoS One 13, e0209577 (2018).

48. Jovanic, T. et al. Competitive Disinhibition Mediates Behavioral Choice and Sequences in Drosophila. Cell 167, 858–870.e19 (2016).

49. Fosque, B. F. et al. Labeling of active neural circuits in vivo with designed calcium integrators. Science (80-.). 347, 755–760 (2015).

50. Briggman, K. L., Abarbanel, H. D. I. & Kristan, W. B. Optical imaging of neuronal populations during decision-making. Science 307, 896–901 (2005).

51. Barker, A. J. & Baier, H. Sensorimotor Decision Making in the Zebrafish Tectum. Curr. Biol. 25, 2804–2814 (2015).

52. Gordus, A., Pokala, N., Levy, S., Flavell, S. W. & Bargmann, C. I. Feedback from Network States Generates Variability in a Probabilistic Olfactory Circuit. Cell 161, 215–227 (2015).

53. Westphal, R. E. & O’Malley, D. M. Fusion of locomotor maneuvers, and improving sensory capabilities, give rise to the flexible homing strikes of juvenile zebrafish. Front. Neural Circuits 7, 108 (2013).

54. Yang, L. et al. Trim9 Regulates Activity-Dependent Fine-Scale Topography in Drosophila. Curr. Biol. 24, 1024–1030 (2014).

55. Yuan, Q. et al. Light-induced structural and functional plasticity in Drosophila larval visual system. Science (80-.). 333, 1458–1462 (2011).

56. Dann, J. F., Buhl, E. H. & Peichl, L. Postnatal dendritic maturation of alpha and beta ganglion cells in cat retina. J. Neurosci. Off. J. Soc. Neurosci. 8, 1485–1499 (1988).

57. Ramoa, A. S., Campbell, G. & Shatz, C. J. Dendritic growth and remodeling of cat retinal ganglion cells during fetal and postnatal development. J. Neurosci. Off. J. Soc. Neurosci. 8, 4239–4261 (1988).

58. Ramoa, A. S., Campbell, G. & Shatz, C. J. Transient morphological features of identified ganglion cells in living fetal and neonatal retina. Science 237, 522–525 (1987).

59. Ren, L., Liang, H., Diao, L. & He, S. Changing dendritic field size of mouse retinal ganglion cells in early postnatal development. Dev. Neurobiol. 70, 397–407 (2010).

60. Couton, L. et al. Development of Connectivity in a Motoneuronal Network in Drosophila Larvae. Curr. Biol. 1–9 (2015). doi:10.1016/j.cub.2014.12.056

61. Tripodi, M., Evers, J. F., Mauss, A., Bate, M. & Landgraf, M. Structural homeostasis: compensatory adjustments of dendritic arbor geometry in response to variations of synaptic input. PLoS Biol. 6, e260 (2008).

62. Yogev, S. & Shen, K. Cellular and Molecular Mechanisms of Synaptic Specificity. 1–21 (2014). doi:10.1146/annurev-cellbio-100913-012953

63. Kumar, V., Zhang, M.-X., Swank, M. W., Kunz, J. & Wu, G.-Y. Regulation of Dendritic Morphogenesis by Ras-PI3K-Akt-mTOR and Ras-MAPK Signaling Pathways. J. Neurosci. 25, 11288–11299 (2005).

64. Threadgill, R., Bobb, K. & Ghosh, A. Regulation of Dendritic Growth and Remodeling by Rho, Rac, and Cdc42. Neuron 19, 625–634 (1997).

65. Sin, W. C., Haas, K., Ruthazer, E. S. & Cline, H. T. Dendrite growth increased by visual activity requires NMDA receptor and Rho GTPases. Nature 419, 475–480 (2002).

66. King, I. & Heberlein, U. Tao kinases as coordinators of actin and microtubule dynamics in developing neurons. Commun. Integr. Biol. 4, 554–6 (2011).

67. Ultanir, S. K. et al. MST3 Kinase Phosphorylates TAO1/2 to Enable Myosin Va Function in Promoting Spine Synapse Development. Neuron 25, 1–15 (2014).

68. Orefice, L. L. et al. Peripheral Mechanosensory Neuron Dysfunction Underlies Tactile and Behavioral Deficits in Mouse Models of ASDs Article Peripheral Mechanosensory Neuron Dysfunction Underlies Tactile and Behavioral Deficits in Mouse Models of ASDs. Cell 1–15 (2016). doi:10.1016/j.cell.2016.05.033

69. Huang, W.-C., Chen, Y. & Page, D. T. Hyperconnectivity of prefrontal cortex to amygdala projections in a mouse model of macrocephaly/autism syndrome. Nat. Commun. 7, 13421 (2016).

70. Ohyama, T. et al. High-Throughput Analysis of Stimulus-Evoked Behaviors in Drosophila Larva Reveals Multiple Modality-Specific Escape Strategies. PLoS One 8, (2013).

71. Tracey Jr., W. D., Wilson, R. I., Laurent, G. & Benzer, S. painless, a Drosophila gene essential for nociception. Cell 113, 261–273 (2003).

72. Han, C., Jan, L. Y. & Jan, Y.-N. N. Enhancer-driven membrane markers for analysis of nonautonomous mechanisms reveal neuron-glia interactions in Drosophila. Proc Natl Acad Sci U S A 108, 9673–9678 (2011).

73. Karuppudurai, T. et al. A Hard-Wired Glutamatergic Circuit Pools and Relays UV Signals to Mediate Spectral Preference in Drosophila. Neuron 81, 603–615 (2014).

74. Groth, A. C., Fish, M., Nusse, R. & Calos, M. P. Construction of transgenic Drosophila by using the site-specific integrase from phage phiC31. Genetics 166, 1775–82 (2004).

75. Hoyer, N., Petersen, M., Tenedini, F. & Soba, P. Assaying Mechanonociceptive Behavior in Drosophila Larvae. BIO-PROTOCOL 8, e2736 (2018).

